# Prolactin action is necessary for parental behavior in male mice

**DOI:** 10.1101/2021.12.30.474525

**Authors:** Kristina O. Smiley, Rosemary S.E. Brown, David R. Grattan

## Abstract

Parental care is critical for successful reproduction in mammals. In comparison to maternal care, the neuroendocrine mechanisms supporting paternal care are less well-studied. Laboratory mice show a mating-induced suppression of infanticide (normally observed in virgins) and onset of paternal behavior. Using this model, we sought to investigate whether the hormone prolactin plays a role in paternal behavior, as it does for maternal behavior. First, using c-fos immunoreactivity in Prlr-IRES-Cre-tdtomato reporter mouse sires, we show that the circuitry activated during paternal interactions contains prolactin-responsive neurons, including the medial preoptic area, bed nucleus of the stria terminalis, and medial amygdala. To evaluate whether prolactin action is required for the establishment and display of paternal behavior, we conditionally deleted the prolactin receptor (Prlr) from 3 distinct cell types: glutamatergic, GABAergic, and CaMKIIα-expressing forebrain neurons. Prlr-deletion from CaMKIIα-expressing forebrain neurons, but not from glutamatergic or GABAergic cells, resulted in a profound effect on paternal behavior, as none of these males completed the pup retrieval task. Finally, although sires do not show an acute increase in circulating prolactin levels in response to pups, pharmacological blockade of prolactin-release at the time of pup exposure resulted in failure to retrieve pups, similar to when the Prlr was deleted from CaMKIIα neurons, with prolactin administration rescuing this behavior. Taken together, our data show that paternal behavior in sires is dependent on basal levels of circulating prolactin acting at the Prlr on CaMKIIα-expressing neurons. These new data in male mice demonstrate that prolactin has a similar action in both sexes to promote parental care.

## INTRODUCTION

Parental care is critical for offspring survival and is an important component of successful reproduction in many species. For mammals, the most common form of parental provisioning is maternal care. However, a number of mammalian species also show paternal care, with males displaying similar parental behaviors as females towards offspring. While less studied than maternal care, many of the brain regions and hormones that regulate maternal care also tend to facilitate paternal behavior. For instance, lesions to olfactory areas, the medial preoptic area (MPOA), and the bed nucleus of the stria terminalis (BNST), which are all crucial to maternal behavior, also disrupt paternal behavior in biparental rodents <u>(reviewed in Bales and Saltzman, 2016; Kohl et al., 2017; Zilkha et al., 2017)</u>. Genetic ablation of galanin-expressing neurons in the MPOA impairs both maternal and paternal care, whereas optogenetic activation of these neurons stimulates parental behaviors in both sexes (Wu et al., 2014). In addition, nonapeptides such as vasopressin (Wang et al., 1994) and oxytocin (Yuan et al., 2019) acting in the brain play a role in promoting paternal behavior, as they do for maternal behavior (Bosch and Neumann, 2012; Bridges, 2020).

While maternal behavior is clearly regulated by the hormonal changes associated with pregnancy (reviewed in Smiley et al., 2019), the role of hormonal signaling in the expression of paternal care has been less clear. We have previously reported that receptors for the anterior pituitary hormone prolactin are expressed throughout the male brain (Kokay et al., 2018), in a pattern similar to that seen in females and particularly in regions known to be important for parental behavior such as the MPOA, BNST, and medial amygdala (MeA). Therefore, we have hypothesized that prolactin action in the brain is important for parental care in males, as it is for females (e.g., Bridges et al., 1990, 2001; <u>Bridges and Mann, 1994; Brown et al., 2017)</u>. While positive correlations between circulating prolactin and paternal care have been observed in a range of mammalian species <u>(e.g., Dixson and Lunn, 1987; Gubernick and Nelson, 1989; Brown et al., 1995; Reburn and Wynne-Edwards, 1999; Carlson et al., 2006)</u>, including humans (Storey et al., 2000; Delahunty et al., 2007; Gettler et al., 2012), not all studies have supported a role for prolactin in paternal care (Brooks et al., 2005; Almond et al., 2006; Wynne-Edwards and Timonin, 2007). We recently showed, however, that species differences in the pattern of prolactin secretion lead to differences in the expression of paternal care. Tight suppression of prolactin secretion by the TIDA neurons in rats is associated with the absence of paternal care, whereas the somewhat less rigid control of prolactin secretion in mice allows for the secretion of higher basal levels of prolactin, which are associated with expression of paternal care (Stagkourakis et al., 2020). Indeed, deletion of Prlr in the MPOA of mice inhibited parental care in male mice (Stagkourakis et al., 2020), as it does in females (Brown et al., 2017). Here, we have used the unique pattern of expression of paternal care in male mice to examine the role of prolactin, investigating the timing of prolactin secretion with respect to the expression of paternal behavior and evaluating the specific regions and neuronal cell types mediating prolactin action on paternal care.

The dramatic change in behavior seen in male ‘father’ mice, creates a valuable model in which the neuroendocrine mechanisms of paternal care can be investigated. Virgin male mice are aggressive towards pups and display infanticidal behaviors or ignore pups (reviewed in Dulac et al., 2014). However, approximately 12 days after mating, infanticidal tendencies are suppressed and males begin to display paternal responses (i.e., pup retrieval) (Vom Saal, 1985). This mating-induced behavioral change requires the act of ejaculation, with no display of paternal responses in animals where only mounting and intromission were allowed (Vom Saal, 1985). The mechanisms behind this remarkable behavioral transition are still unknown. In females, prolactin (or its placental homologue, placental lactogen) is high during pregnancy, and further stimulated by the suckling stimulus during interactions with pups (Phillipps et al., 2020). The pattern of prolactin secretion during the expression of paternal care, however, is much less apparent. It is possible that interactions of the males with pups induces prolactin secretion, promoting paternal care. Alternatively, as ejaculation causes a significant rise in circulating prolactin in both humans and rodents (Krüger et al., 2002; Valente et al., 2021), another possibility is that this ejaculation-induced surge in prolactin secretion is involved in the transition away from infanticidal behavior and the subsequent expression of paternal care.

To test these hypotheses, we have firstly investigated whether prolactin-responsive neurons are activated by exposure to pups. We found that paternal interactions in fathers evoke a pattern of c-fos activity in prolactin-responsive neurons across multiple brain regions that is not present in virgins that are exposed to pups and do not exhibit paternal behavior. We then used a combination of conditional prolactin receptor (Prlr) deletion in specific populations of neurons and pharmacological suppression of prolactin to investigate the role of endogenous prolactin signaling in the expression of paternal care. We found that either a Prlr knockout in CaMKIIα-expressing neurons (but not deletion of Prlr in either GABAergic or glutamatergic neurons) or suppression of circulating prolactin specifically during exposure to pups markedly disrupted paternal care. In contrast, suppression of the mating-induced release of prolactin had no effect on the transition away from infanticide and did not significantly alter the subsequent expression of paternal care. Taken together, these data provide substantial evidence that prolactin action in the brain at the time of exposure to pups is necessary for the display of normal paternal behaviors, likely mediated through multiple different populations of neurons in the paternal circuit.

## RESULTS

### Prolactin-responsive neurons are activated by pup exposure in sires

Our first aim was to identify whether the neural circuitry activated during paternal interactions contained prolactin-responsive neurons. To address this, we measured c-fos immunoreactivity (a marker of recent neural activation) in Prlr-IRES-Cre-tdtomato reporter mouse sires that were either exposed to pups using a standard pup exposure assay (Fig 1A), or received no pup exposure (controls). Latency to retrieve all 4 pups into the nest, and durations of paternal behaviors including sniffing, retrieving, and huddling pups, as well as time spent nesting alone were quantified for each male, with all animals retrieving the pups to a nest and huddling over pups for the majority of the test (Fig 1B). Consistent with previous reports, the ventral part of the bed nucleus of the stria terminalis (BSTv), posteroventral division of the medial amygdaloid nucleus (MeApd), and medial preoptic area (MPOA) showed pup-induced increases in c-fos immunoreactivity, relative to non-exposed control animals (Fig 1K,T, CC; Table 1). A significant portion of prolactin-responsive cells were activated in response to paternal interactions, with pup-exposed fathers showing a 2-4 fold increase in c-fos expression in Prlr-expressing cells in the BSTv (Fig 1L) and MPOA (Fig 1DD), with the MeApd (Fig 1U) showing an 12-fold increase over the very low levels of c-fos seen in controls (Table 1). In comparison, pup-exposed Prlr-IRES-Cre-tdtomato virgin males, which are generally aggressive towards pups, showed significant pup exposure-induced c-fos immunoreactivity in Prlr-containing cells in the MeApd and periventricular nucleus (PVN) (Fig 2V, NN; Table 2). However, unlike sires, there was no significant pup-induced c-fos in Prlr-containing cells in the MPOA or BSTv in virgin males (Fig 2M,EE; Table 2) or in recently mated males (Fig S1I,R; Table S1). Together these data show that although Prlr-expressing neurons in the MeApd respond to pups in both virgins and sires, activation of Prlr-expressing neurons in the MPOA and BSTv only occurs after pup-interactions in sires.

**Figure 1.**
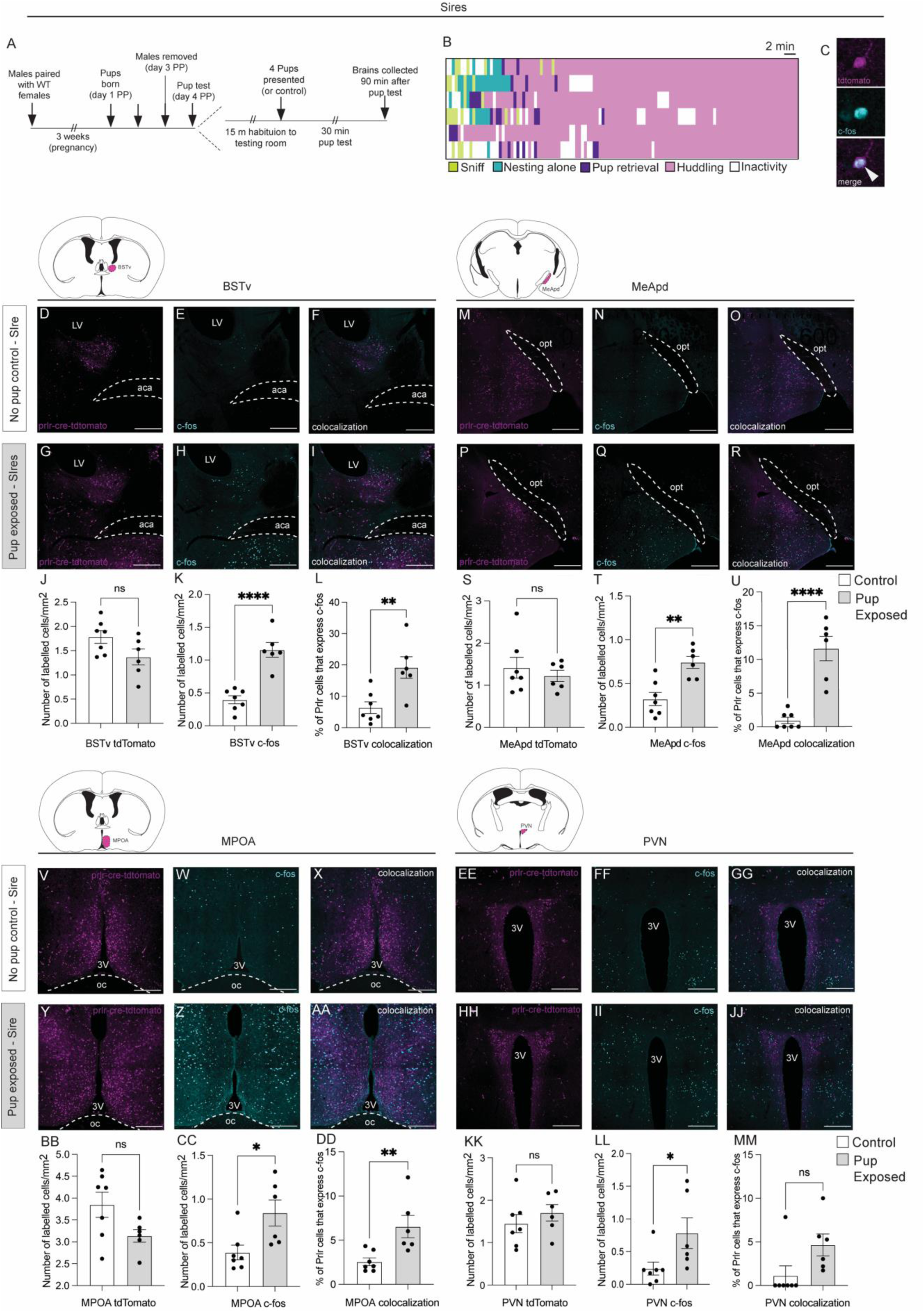
Prolactin-responsive neurons are activated by pup exposure in sires. Male Prlr-IRES-Cre/tdtomato reporter mice were used to identify pup-induced cell activation (c-fos) in prolactin receptor (Prlr)-containing neurons. (A) Schematic of pup retrieval test used to induce c-fos in response to pup interactions. WT= wildtype; PP=post-partum. (B) Ethogram of paternal behavior during the 30 min pup test. Note that each pup-exposed male (n=6) retrieved pups to the nest and spent the majority of the time huddling over pups. (C) Representative high-powered images showing a tdtomato (Prlr) labelled cell (magenta), c-fos immunoreactivity (cyan), and the merged image to show an example of a colocalized cell (white arrow). (D-R, V-JJ) Representative images of tdtomato labelling (indicative of Prlr; magenta), c-fos labelling (cyan), and colocalization of tdtomato+c-fos (white) in control (top rows) and pup-exposed sires (bottom rows). The brain regions examined are indicated by the colored areas on atlas drawings; ventral part of the bed nucleus of the stria terminalis (BSTv), posteroventral division of the medial amygdaloid nucleus (MeApd), medial preoptic area (MPOA), and periventricular nucleus (PVN). (L,U,DD) Pup-exposed males (grey bars) had significantly higher c-fos expression in Prlr-expressing cells in the BSTv, MeApd, and MPOA compared to control males (white bars). Bar graphs show individual data points (black circles) and mean ± SEM; ns = non-significant (p>0.05), *p<0.05, **p<0.01, ***p<0.001,**** p<0.0001. 3V = third ventricle, aca = anterior commissure, oc = optic chiasm, opt = optic tract. Scale bars = 50 μm.

**Figure 2.**
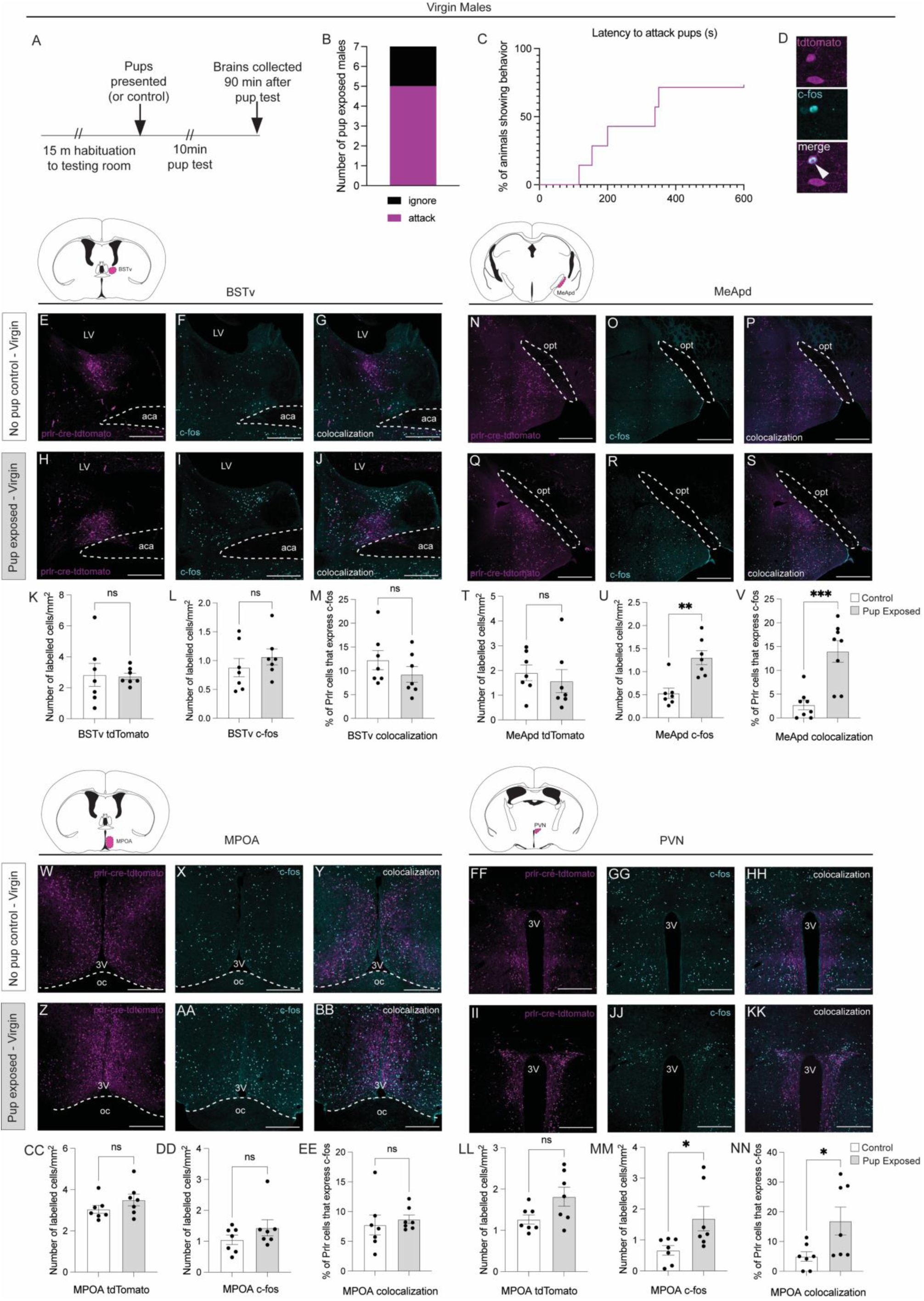
Prolactin-receptor containing neurons in the PVN and MeApd are activated by pup exposure in virgin mice. (A) Schematic of pup exposure assay used with virgin Prlr-IRES-Cre-tdtomato males (n=7 per group). (B) Five out of the 7 pup exposed males tested attacked pups, while 2 males ignore pups. (C) Latency to attack pups across all males. Note that males were only exposed to pups for 10 min or until attacking occurred, in which case, pups were immediately removed and testing ceased. (D) Representative high-powered image showing a tdtomato/Prlr labelled cell (magenta), c-fos immunoreactivity (cyan), and the merge of the two images to show an example of a colocalized cell (white arrow). (E-S, W-KK) Representative images of tdtomato labelling (indicative of Prlr; magenta), c-fos labelling (cyan), and colocalization of tdtomato+c-fos are shown for each brain region examined in control virgins (top rows) and pup-exposed virgins (bottom rows). The brain regions examined are indicated by the colored areas on atlas drawings; ventral part of the bed nucleus of the stria terminalis (BSTv), posteroventral division of the medial amygdaloid nucleus (MeApd), medial preoptic area (MPOA), and periventricular nucleus (PVN). Note that pup-exposed virgin males (grey bars) had significantly more c-fos expression in Prlr-expressing cells compared to control males (white bars) in the MeApd and PVN, but not the BSTv or MPOA. Bar graphs show individual data points (black circles) and mean ± SEM; (s)=seconds; ns = non-significant (p>0.05), *p<0.05, **p<0.01, ***p<0.001. 3V = third ventricle, aca = anterior commissure, oc = optic chiasm, opt = optic tract. Scale bars = 50 μm.

**Table 1.**
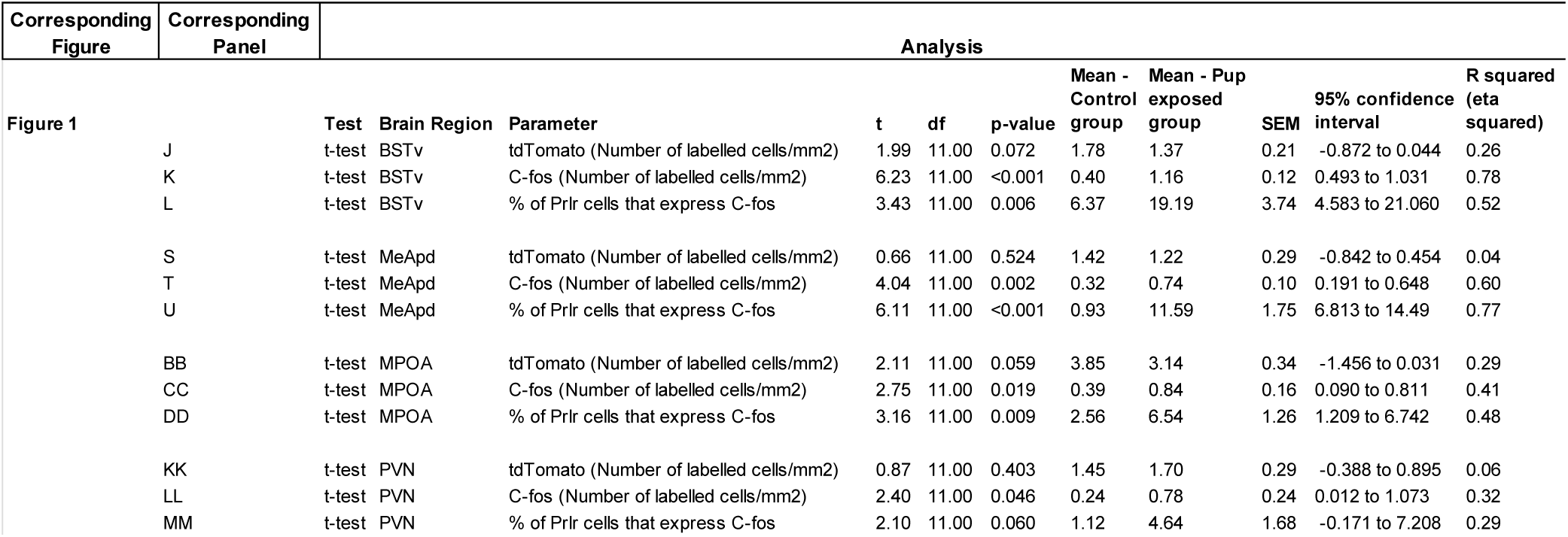
Statistical analysis for Figure 1

**Table 2.**
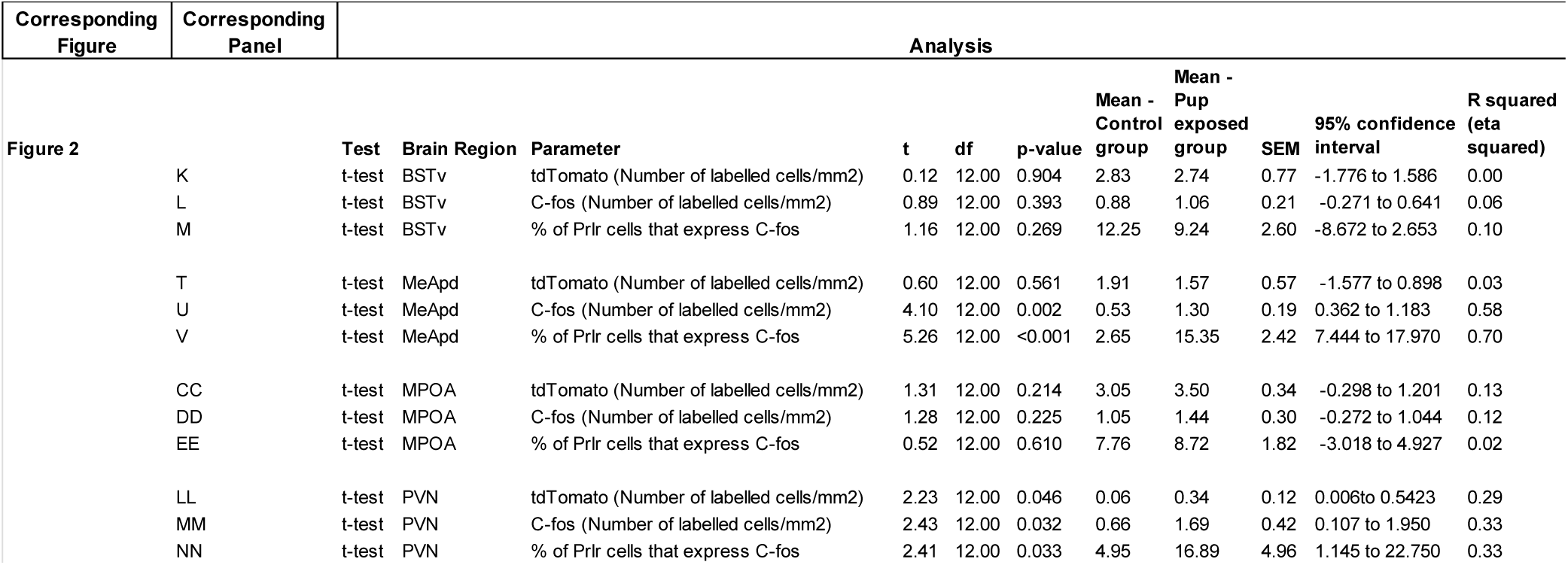
Statistical analysis for Figure 2

### Prolactin receptor knockout in forebrain CaMKIIα-expressing neurons impairs paternal behavior

Following identification that populations of Prlr-expressing cells exhibited pup-induced c-fos immunoreactivity (Fig 1), we aimed to investigate (1) whether the Prlr was necessary for paternal behavior in mouse sires and, if so, (2) which population of Prlr-expressing cells were necessary. Using the cre-lox strategy, we generated 3 different conditional Prlr knockout lines which targeted either glutamatergic neurons (Prlr Vglut KO), GABAergic neurons (Prlr Vgat KO), or CaMKIIα expressing-neurons (which targets predominantly a population of forebrain neurons including some excitatory and inhibitory neurons; Benson et al., 1992; Liu and Murray, 2012; Prlr CKC KO). The degree and distribution of Prlr deletion was assessed across the forebrain by eGFP immunoreactivity, which is induced upon Cre-mediated recombination of the transgene (Brown et al., 2016a), with respective littermate Cre-negative Prlr^lox/lox^ control mice (with intact *Prlr*) showing no eGFP expression (Fig 3; Table 3). Note that in all models there was some Prlr deletion in every brain region examined, but the degree of this deletion was markedly different in each mouse line, reflecting the composition of neuronal sub-types expressed in each region.

**Figure 3.**
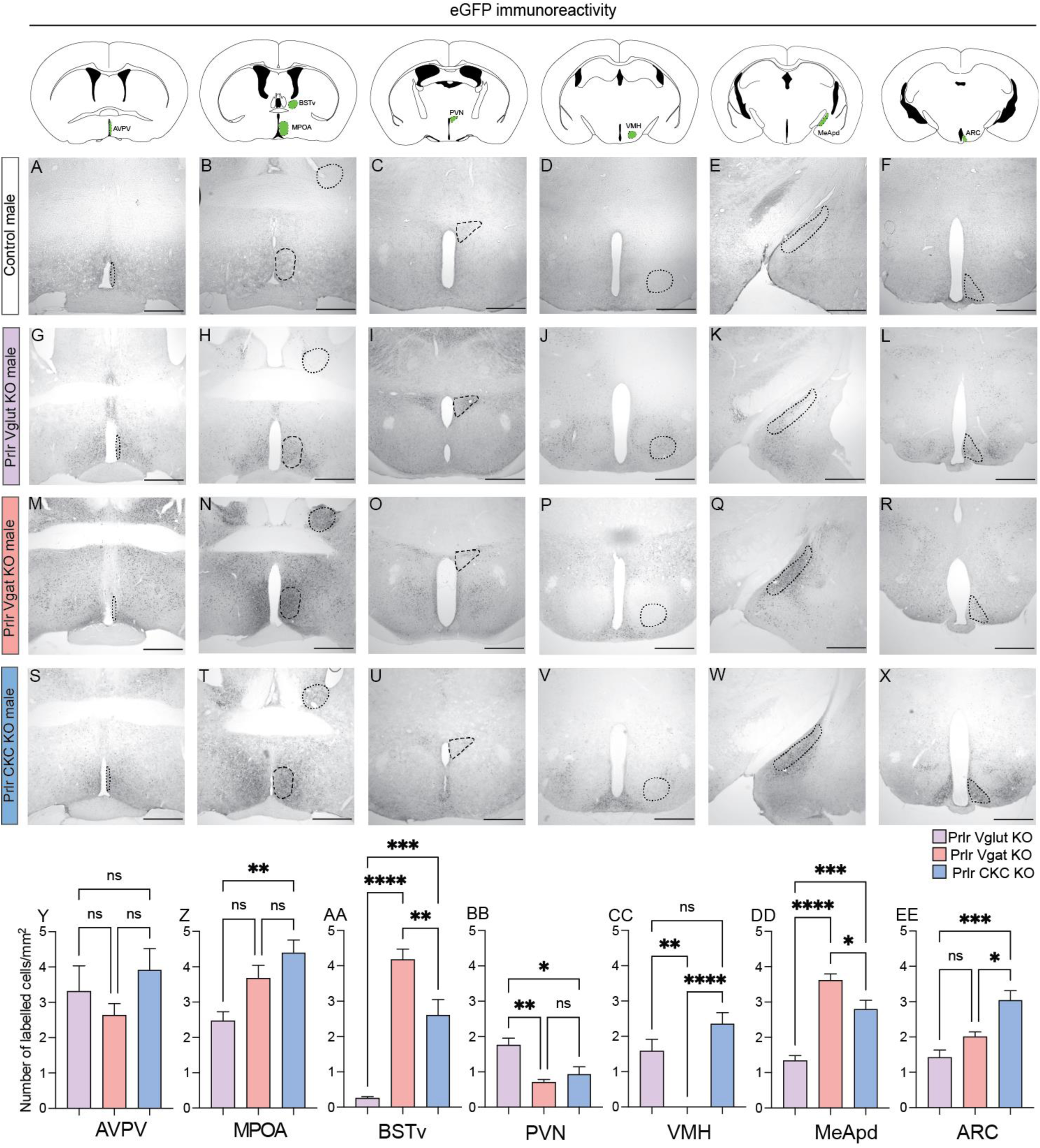
Varying degrees of prolactin receptor deletion are observed in Prlr Vglut, Vgat, and CKC KO males. The Prlr^lox/lox^ mouse model used in this paper was designed such that Cre-mediated inversion deletes the Prlr gene and knocks eGFP into place, meaning eGFP can be used as marker for successful recombination (e.g., Prlr deletion). We compared eGFP immunoreactivity across the forebrain of male Prlr^lox/lox^/VGlut-Cre+ (Prlr Vglut KO, purple bars), Prlr^lox/lox^/VGat-Cre+ (Prlr Vgat KO, orange bars), and Prlr^lox/lox^/CamK-Cre+ (Prlr CKC KO, blue bars) (n=6 per group). Colored areas on atlas drawings indicate the brain region that was quantified. Note that none of control brains from any group showed eGFP expression in any part of the brain (indicative of no recombination/Prlr deletion; e.g., intact *Prlr*), so only one set of control brains (Prlr CKC Cre-negative) is shown for comparison (A-F), and were not included in the analysis. Representative images from Prlr Vglut KO (G-L), Prlr Vgat KO (M-R), and Prlr CKC KO (S-X) brains. Note that in all models there was some Prlr deletion in every brain region examined, but the degree of this was markedly different in each mouse line, reflecting the composition of neuronal sub-types expressed in each region. Bar graphs are presented as mean ± SEM; ns = non-significant (p>0.05), *p<0.05, **p<0.01, ***p<0.001,**** p<0.0001. AVPV= anteroventral periventricular nucleus, MPOA= medial preoptic area, BSTv= ventral part of the bed nucleus of the stria terminalis, PVN= periventricular nucleus, VMH=ventromedial hypothalamus; MeApd= posteroventral division of the medial amygdaloid nucleus; ARC=arcuate nucleus. Scale bars = 50 μm.

**Table 3.**
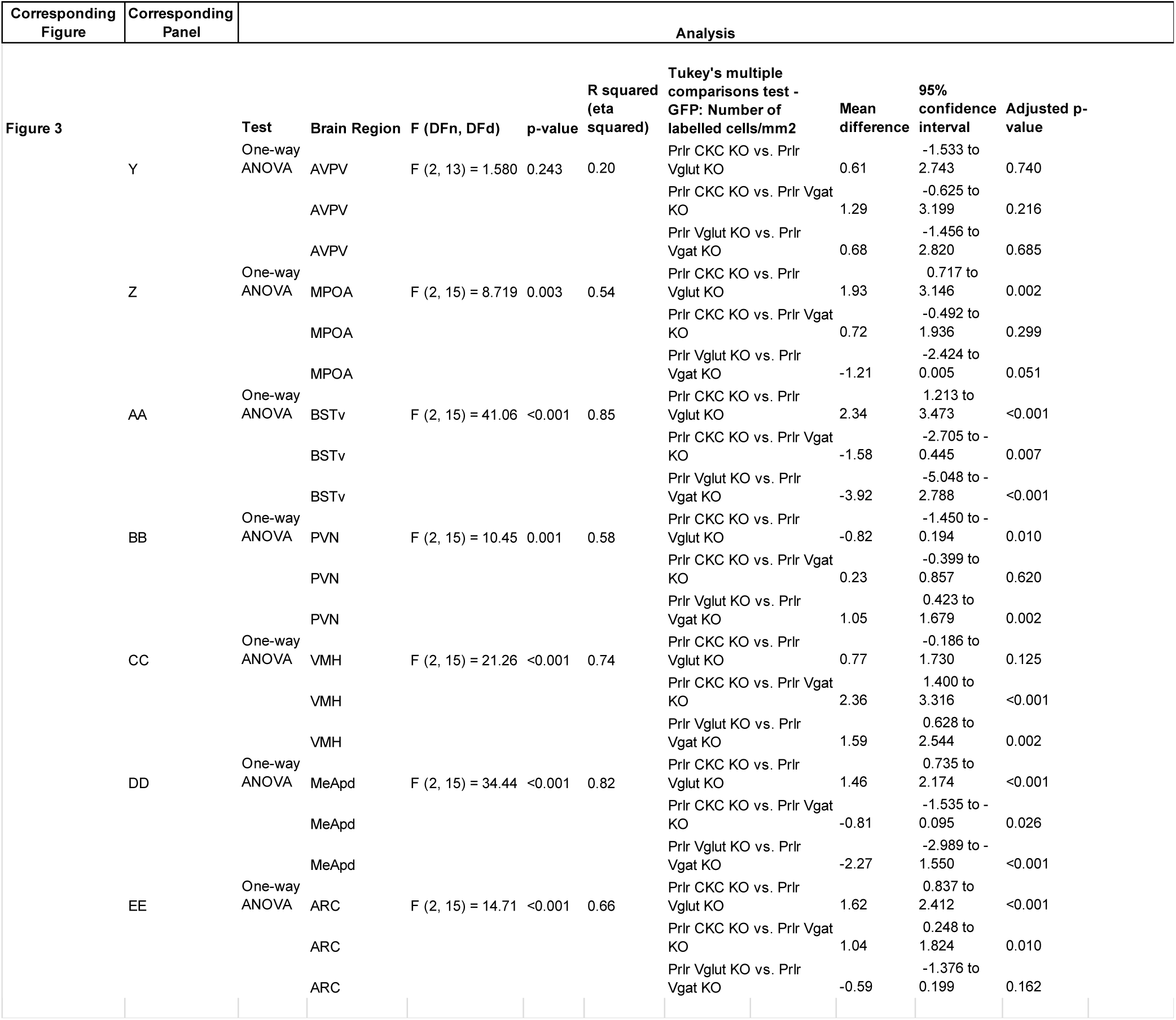
Statistical analyses for Figure 3

Adult males from all 3 lines were tested as sires for paternal behavior using the pup retrieval test (Fig 1A). Neither Prlr Vglut KO or Prlr Vgat KO males showed any detectable deficits in paternal behaviors compared to Cre-negative control males (Fig 4A,B, D-O, Table 4). In contrast, we observed significant deficits in Prlr CKC KO males, as none of these males retrieved all pups to the nest (Fig 4C,P,Q; Table 4; Movie S1). Although not statistically different, only 2 of the 7 Prlr CKC KO males displayed normal huddling behavior (Fig 4T), with most males spending the majority of the time in the nest alone, without pups (Fig 4U), suggesting additional paternal behaviors beyond retrieval were also impaired. Prlr CKC KO males showed a marked reduction in functional Prlr as assessed by pSTAT5 (a robust marker for Prlr activation (Brown et al., 2010); Fig 5; Table 5) in a number of forebrain regions that correspond with Prlr deletion (Fig 3S-X) including the anteroventral periventricular nucleus (AVPV), MPOA, ventromedial hypothalamus (VMH), MeApd, and the arcuate nucleus (ARC). These results indicate that Prlr expression in CaMKIIα expressing-neurons is necessary for paternal care behavior and that multiple regions may be involved in a prolactin-sensitive network that controls pup retrieval behavior.

**Figure 4.**
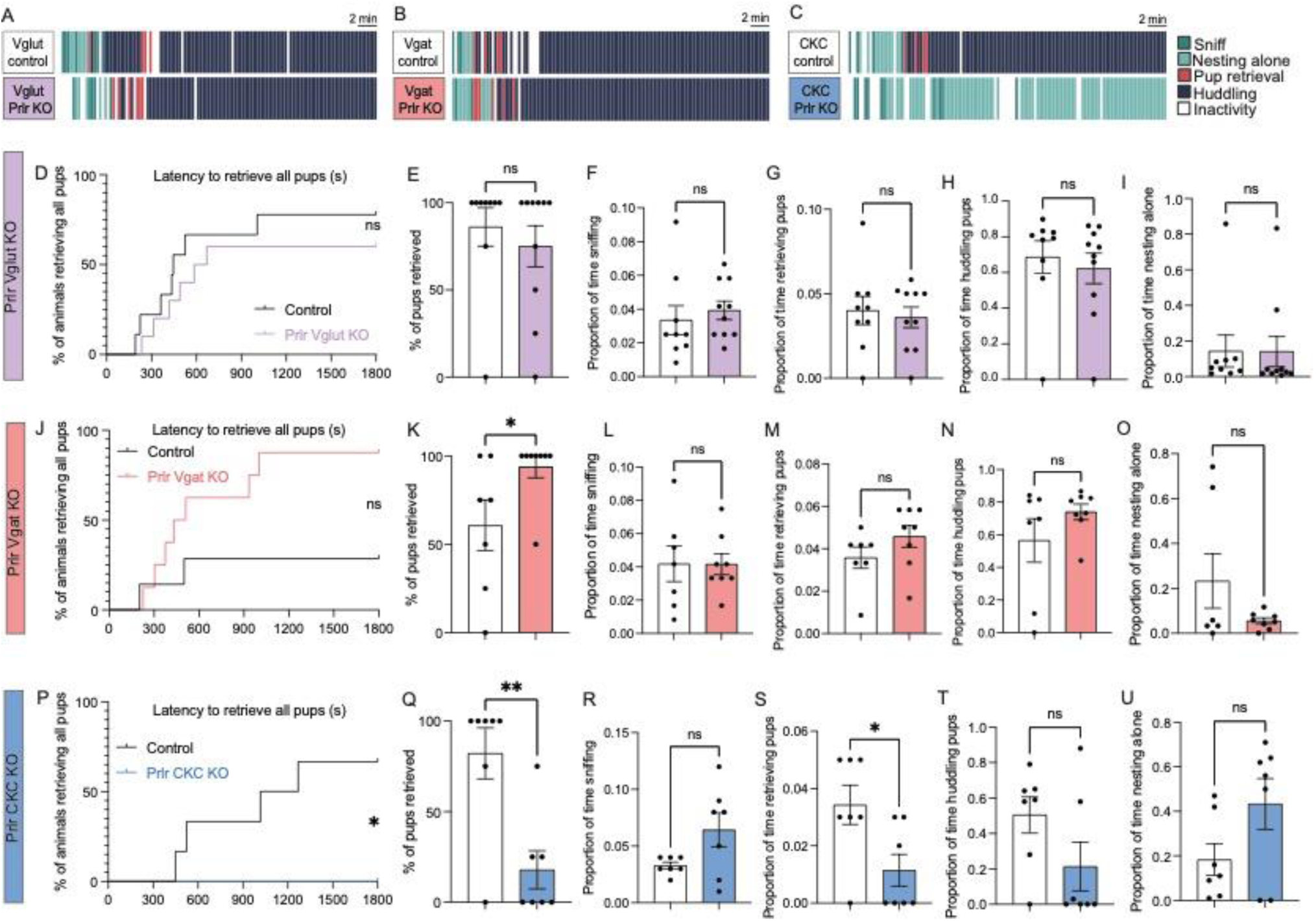
Prolactin receptor knockout in CaMKIIα-expressing forebrain neurons disrupts paternal behavior. (A-C) Representative ethograms of paternal behavior observed during the pup retrieval test in Prlr^lox/lox^/VGlut-Cre+ (Prlr Vglut KO), Prlr^lox/lox^/VGat-Cre+ (Prlr Vgat KO), and Prlr^lox/lox^/CamK-Cre+ (Prlr CKC KO) sires compared to respective Cre-negative Prlr^lox/lox^ control sires. (D-O) Prolactin-receptor knockout in glutamergic neurons (purple bars, n=10) or GABAergic (orange bars, n=8) did not result in any deficits in paternal behavior, compared to control males (white bars, n=9, 7 respectively). (P-U) In contrast, Prlr CKC KO males showed significant deficits in pup retrieval behaviors, with none of the Prlr CKC KO males retrieving the full set of 4 pups to the nest (P,Q). Note that although not statistically different from controls, most Prlr CKC KO males spent the majority of their time nesting alone and not huddling pups (T,U). For all bar graphs, data points represent individual subjects and are presented as mean ± SEM; ns = non-significant (p>0.05), *p<0.05, **p<0.01.

**Figure 5.**
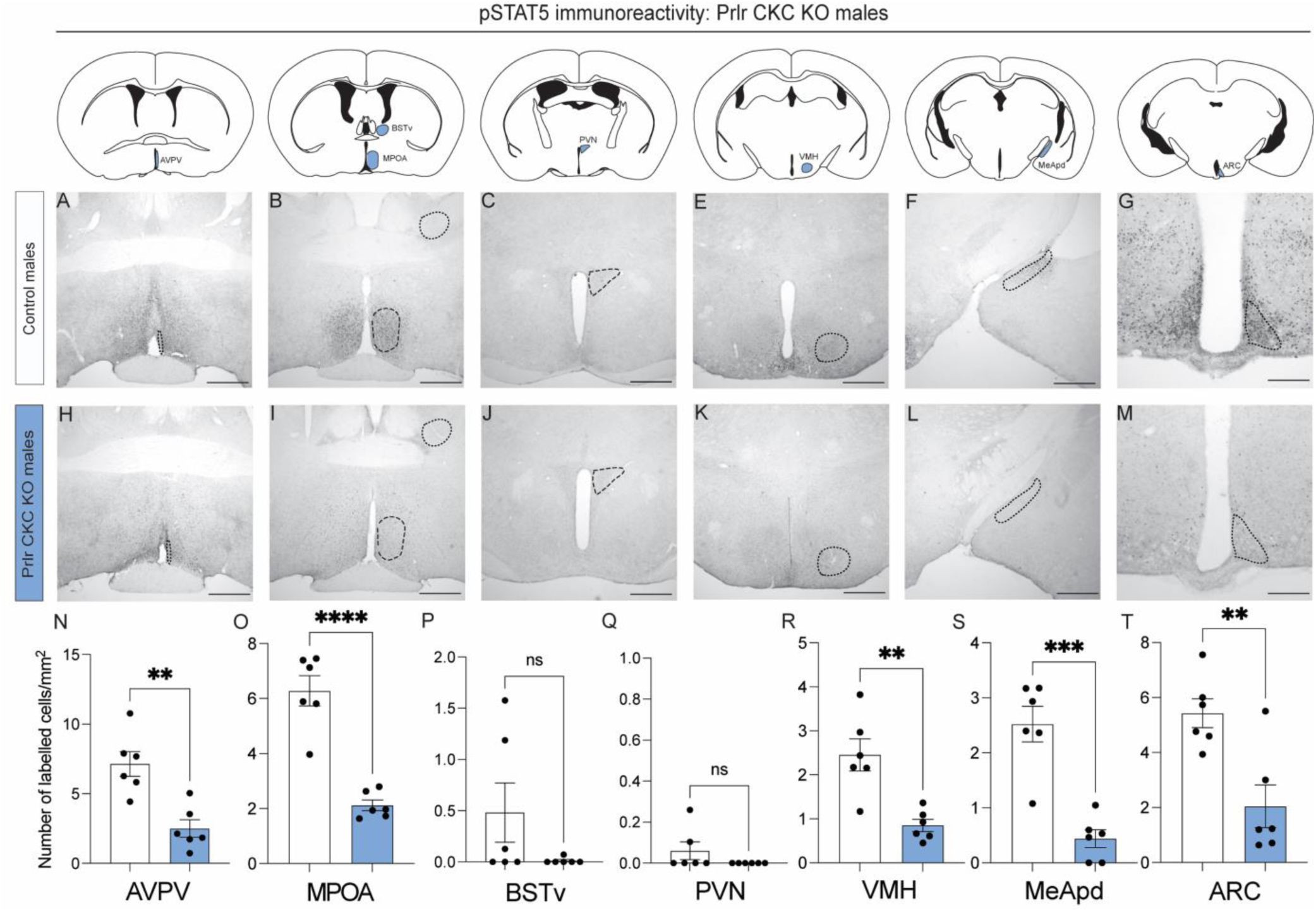
pSTAT5 activity is significantly reduced in Prlr CKC KO males. (A-M) Prlr CKC KO males (n=6) and cre-negative control males (n=6) were injected with ovine prolactin 45 min before perfusion to assess pSTAT5 (a reliable marker of Prlr activation) immunoreactivity across the forebrain. Colored areas on atlas drawings indicate the brain regions that were quantified. Note that control males (with fully intact *Prlr*) show a robust response to prolactin (high levels of pSTAT5 immunoreactivity) (A-G), whereas Prlr CKC KO males show markedly reduced levels of pSTAT5 activity in response to prolactin (H-M) confirming significant Prlr deletion in the AVPA, MPOA, VMH, MeApd, and ARC (N-T). AVPV= anteroventral periventricular nucleus, MPOA= medial preoptic area, BSTv= ventral part of the bed nucleus of the stria terminalis, PVN= periventricular nucleus, VMH=ventromedial hypothalamus; MeApd= posteroventral division of the medial amygdaloid nucleus; ARC=arcuate nucleus. For all bar graphs, data points represent individual subjects and are presented as mean ± SEM; ns = non-significant (p>0.05), **p<0.01, ***p<0.001,**** p<0.0001. Scale bars = 50 μm.

**Table 4.**
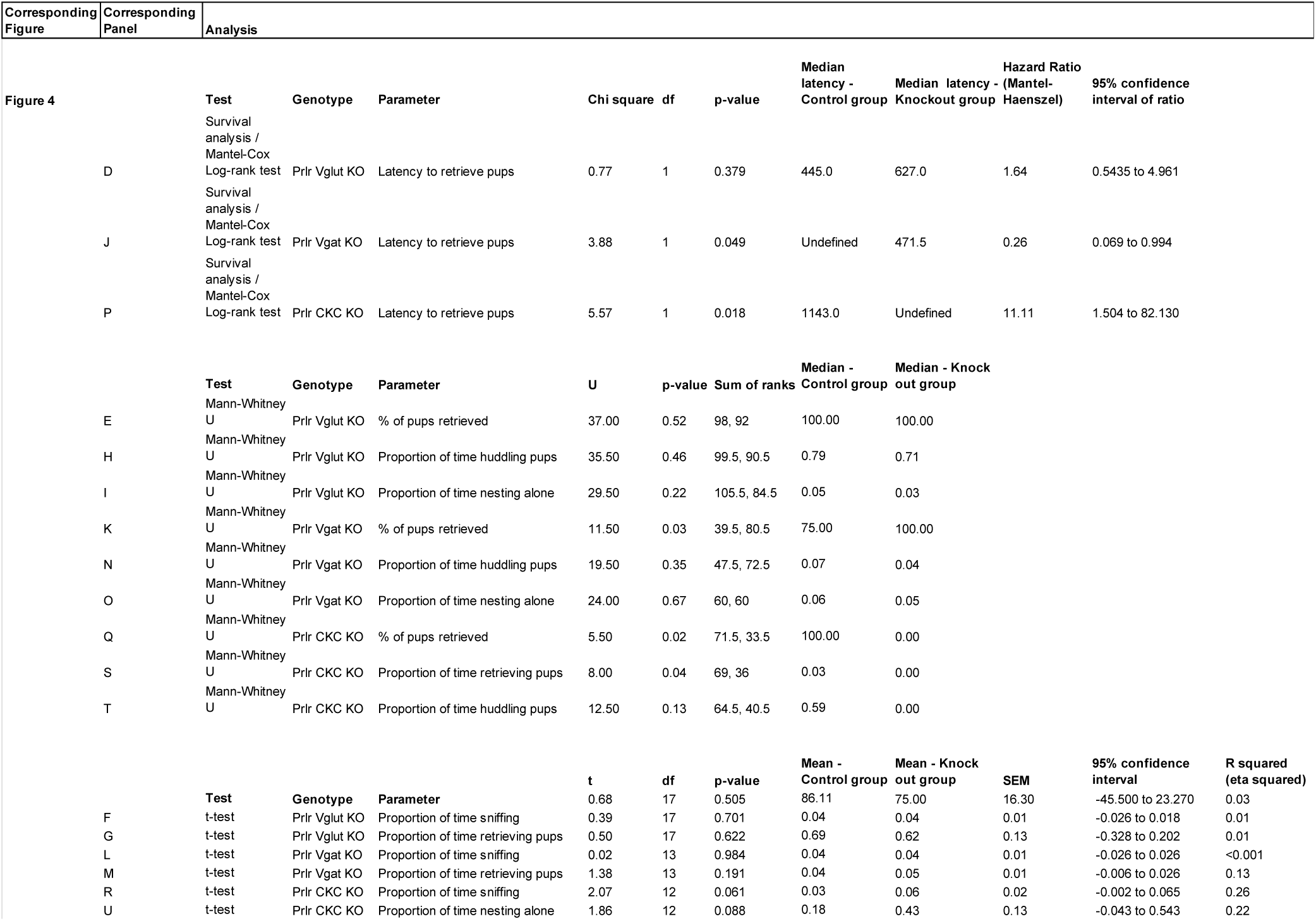
Statistical analyses for Figure 4

**Table 5.**
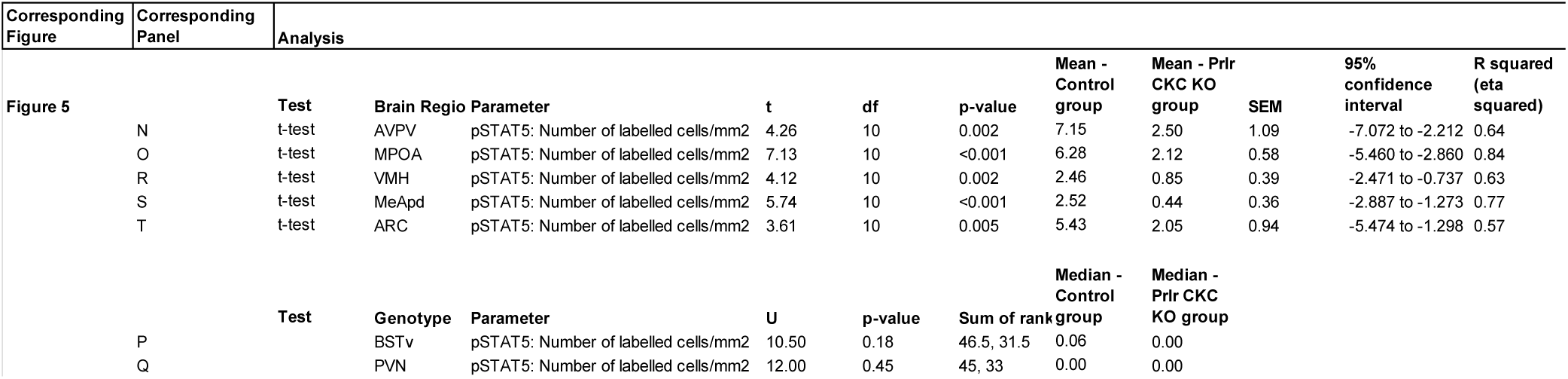
Statistical analyses for Figure 5

### Circulating prolactin is required for paternal behavior in sires

With Prlr expression in the brain clearly necessary for paternal behavior, we subsequently aimed to investigate the temporal dynamics of circulating prolactin to identify critical periods of prolactin exposure for paternal behavior. Virgin males have relatively low prolactin levels, which do not differ between the light and dark cycles (Fig 6A; Table 6). Consistent with previous reports (Valente et al., 2021), males showed a transient mating-induced surge of prolactin that lasted 30 min post-ejaculation (Fig 6B; Table 6). Surprisingly, we saw no increase from circulating prolactin from virgin levels in sires during the pup rearing period (Fig 6C; Table 6).

**Figure 6.**
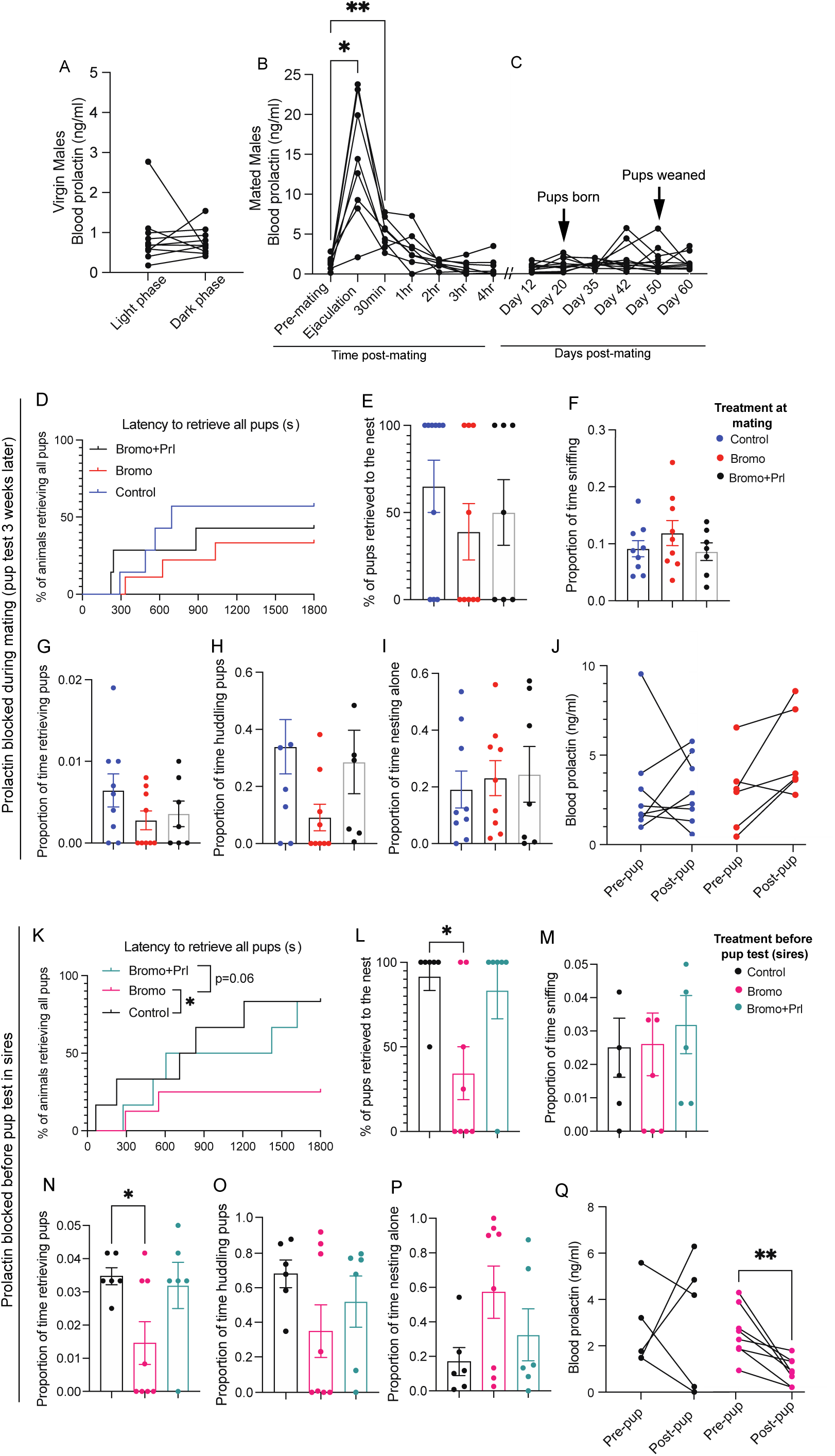
Circulating prolactin is required to show paternal behaviors in sires. (A-C) Prolactin concentrations measured in blood samples taken from C57BL/6J male mice. Virgin male mice have relatively low prolactin levels, which do not differ between the light and dark cycles (A), but show a transient mating-induced surge of prolactin that lasts up to 30 min post-ejaculation (B) Asterisks indicate when prolactin is significantly higher than pre-mating baseline concentrations. In contrast, we did not detect an increase in prolactin in sires during the pup rearing period (C). (D-J) C57BL/6J male mice were treated with either bromocriptine (to prevent prolactin secretion from the pituitary; n=9) or vehicle-control (n=7) 1.5 hrs prior, or bromocriptine 1.5 hours and ovine prolactin 45 min prior to their first mating experience. Paternal behaviors were assessed 3 weeks later (the normal length between mating and birth of pups) using a pup retrieval assay. Neither bromocriptine or bromocriptine+prolactin had any effect on the pup retrieval latency (D), the number of pups retrieved (E) or any paternal behaviors measured (F-I). Prolactin levels were measured before and after the 30 min pup test and were not affected by previous bromocriptine treatment at mating (J). (K-Q) C57BL/6J sires were treated with either bromocriptine (n=8) or vehicle (n=6) 1.5 hrs before or bromocriptine 1.5 hours and ovine prolactin 45 min before the pup retrieval test. Bromocriptine-treated sires retrieved significantly less pups compared to controls (K,L). Other behaviors including sniffing (M), huddling (O), and nesting alone (P) were not statistically different. Bromocriptine+prolactin treated males were not significantly different from controls. Note that pup interactions did not increase prolactin in vehicle-control males (Q). Bar graphs show individual data points (circles) and mean ± SEM; ns = non-significant (p>0.05), *p<0.05, **p<0.01.

**Table 6.**
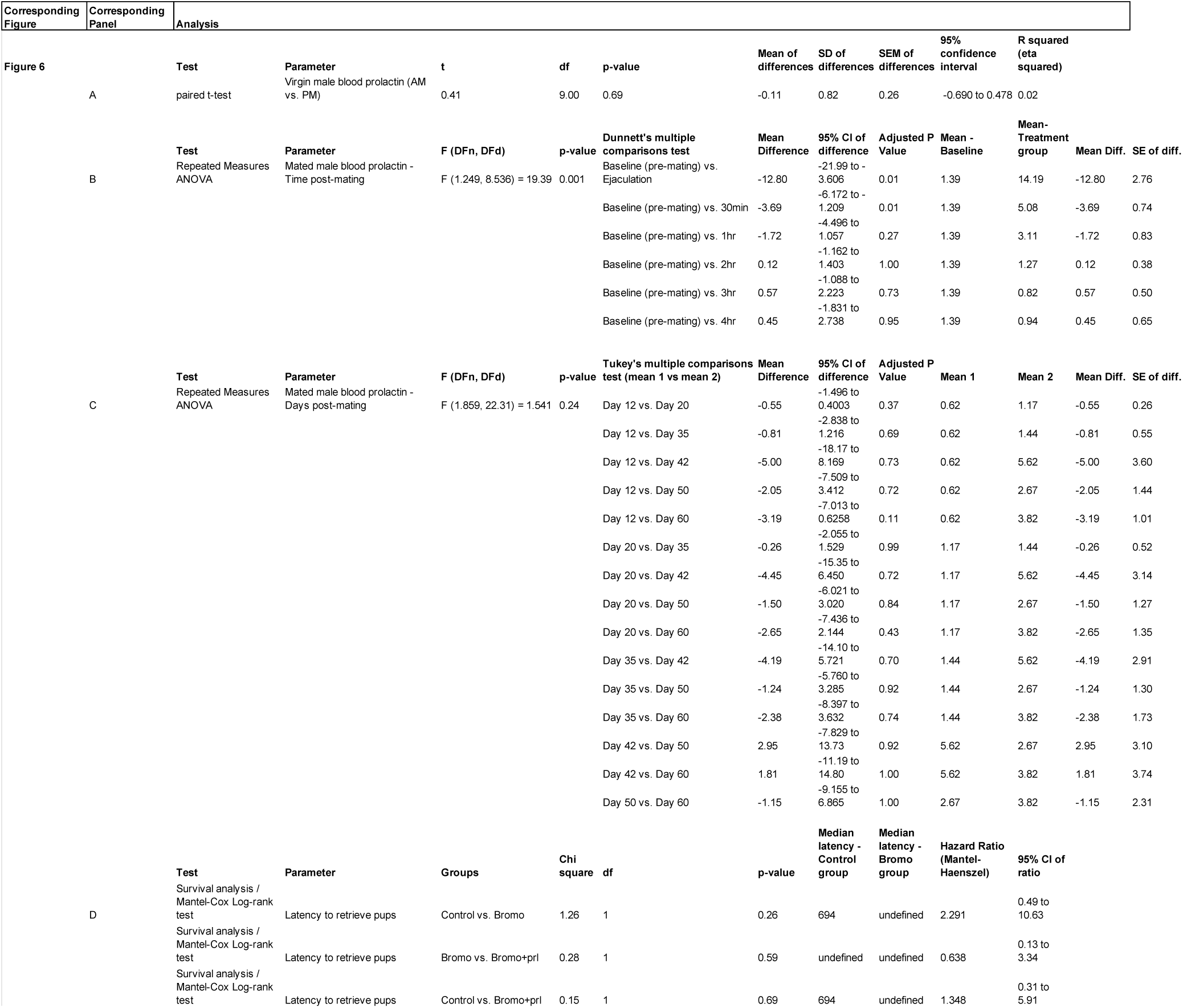
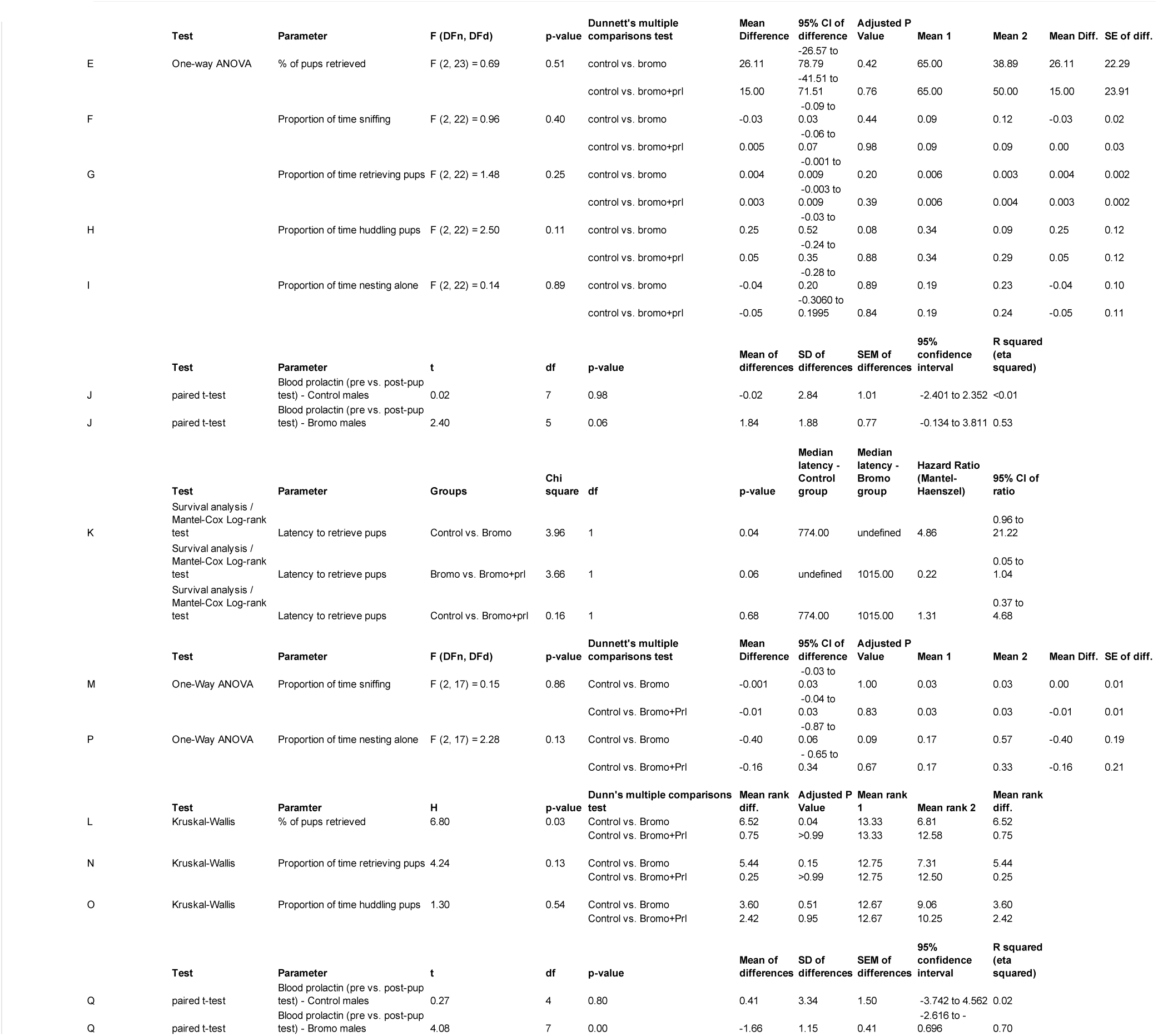
Statistical analyses for Figure 6

As paternal care in laboratory mice is dependent on ejaculation during mating (Vom Saal, 1985), we hypothesized that mating-induced prolactin may be required to signal the transition into paternal care. However, preventing the mating-induced prolactin surge with bromocriptine (Fig S2A), a well-established prolactin inhibitor (Brown et al., 2010) which does not affect mating behavior (Fig S2B-F; Table S2), had no significant effects on subsequent paternal behavior or prolactin levels when tested 3 weeks later (the normal time between mating and birth of offspring; Fig 6D-J; Table 6). Likewise, administering both bromocriptine and prolactin (rescue control) at mating did not significantly affect subsequent paternal behavior. Notably, bromocriptine-treated animals did not show any infanticidal responses, indicating that mating-induced prolactin release is not required for the transition to paternal care, nor the suppression of infanticidal behaviors.

To determine whether paternal behavior in sires is dependent on circulating prolactin at the time of pup care, we next administered bromocriptine, bromocriptine+prolactin (rescue control), or vehicle to mouse sires prior to the pup retrieval test (Fig 1A). Accompanying the suppression of circulating prolactin (Fig 6Q; Table 6) were significant deficits in pup retrieval behavior, similar to that seen in the Prlr CKC KO males, with most males failing to retrieve all of the pups during the task (Fig 6K, L; Table 6). Notably, administering prolactin following bromocriptine rescued this pup retrieval behavior, with nearly all bromocriptine+prolactin males retrieving pups, similar to controls. Although not statistically different, most bromocriptine-treated males spent the majority of the test nesting alone without pups, and only 3 out of the 7 bromocriptine males displayed huddling behavior within the range of control huddling behavior (Fig 6O). Pup interactions did not cause an acute increase in prolactin in control fathers (Fig 6Q; Table 6). Note that as above, no males showed any infanticidal responses. This effect of prolactin is unique to sires as prolactin exposure did not affect virgin male behavior (Fig S3; Table S3). Together, these data demonstrate that prolactin action is specifically required at the time of pup exposure in father mice for the display of paternal responses.

## DISCUSSION

Our data indicate that prolactin action is necessary for the normal expression of paternal care in laboratory mice. Specifically, we found that (1) paternal interactions with pups induces c-fos immunoreactivity in prolactin-responsive neurons in multiple regions including the BSTv, MPOA, and MeApd in fathers, (2) males with a CaMKIIα-specific deletion of the Prlr displayed significant deficits in pup retrieval behavior, and (3) these effects are driven by basal circulating prolactin, which is required at the time of pup care in sires for the normal display of paternal behavior, even though there is no specific activation of prolactin secretion in response to pups. Taken together, these studies provide strong causal evidence that prolactin acting on the Prlr in the brain is required for the normal paternal responses in mice, providing a long-sought explanation for a physiological role for this hormone in a male mammal.

As a first step to determine whether prolactin is involved in mouse paternal behavior, we measured c-fos activity in response to pup interactions in the BSTv, MPOA, MeApd, and PVN of father mice (which retrieve pups) and virgin male mice (which attack pups). While this is not an exhaustive list of areas that could be potentially involved in paternal care, roles for these regions have been established in other paternal mammals (<u>reviewed in Bales and Saltzman, 2016; Kohl et al., 2017</u>; <u>Zilkha et al., 2017</u>) *and* express the Prlr (Kokay et al., 2018). Using this method, we found that Prlr expressing cells were activated in response to pups in the MPOA and BSTv of fathers and in the MeApd of both fathers and virgins. The medial amygdala (MeA) is involved in downstream processing of sensory cues controlling social behavior and is highly connected with the MPOA and BST (Numan and Insel, 2003; Kohl et al., 2017). GABAergic neurons in the MeA have been shown to regulate both infanticidal and parenting behaviors in male mice in an activity level dependent manner (Chen et al., 2019), but the role of prolactin in the MeApd during parenting has yet to be explored. The MPOA and BST are highly interconnected, mainly through GABAergic connections (Tachikawa et al., 2013), and have long been known to be critical nodes in the parenting behavior circuits (Numan and Insel, 2003). Prlr-containing neurons in the MPOA have been shown to be required for both maternal (Brown et al., 2017) and paternal (Stagkourakis et al., 2020) behaviors, however, the role of prolactin/Prlr in the BST is less studied. These areas were not significantly activated in virgin or mated males, indicating pup stimuli elicits neuronal activity in Prlr-expressing cells in the MPOA and BSTv specifically in fathers, which may depend on prior mating experience and/or other internal factors, including other hormonal changes, that co-occur in fathers.

Following identification that populations of Prlr-expressing cells exhibited pup-induced c-fos immunoreactivity in fathers, we aimed to determine whether the Prlr in these circuits were necessary for paternal behavior. Given the widespread distribution of c-fos-responsive Prlr neurons, we focused on three broad subpopulations of neurons found throughout these regions: Glutamate, GABA, and CaMKIIα. Males with Prlrs genetically deleted from either glutamatergic (Prlr Vglut KO) or GABAergic (Prlr Vgat KO) cells did not show any deficits in paternal behavior. Similar results have been found in female mice, where a Prlr Vglut KO, Prlr Vgat KO, or combined Prlr Vglut+Vgat KO did not result in any significant deficits in maternal behavior, whereas complete deletion of Prlr from all neurons in the MPOA did profoundly affect maternal behavior (Brown et al., 2017). In males, we found that deleting the Prlr out of CaMKIIα-expressing forebrain neurons (Prlr CKC KO), which would include multiple different neuronal cell types, resulted in a profound effect on paternal behavior, with none of the Prlr CKC KO males retrieving the full set of pups to the nest. Although recombination in the CaMKIIα-Prlr ^lox/lox^-Cre-line is restricted to post-mitotic neurons, one consideration of these findings is that other postnatal developmental effects resulting from Prlr deletion could account for some of the effects observed. However, this is unlikely as the pharmacological blockade of circulating prolactin manipulations in fathers were undertaken in adulthood and reproduced equivalent deficits in pup retrieval behavior. Considering earlier findings that Prlr-containing neurons in the MPOA are critical for paternal care in mice (Stagkourakis et al., 2020), it is possible that a specific subpopulation of CaMKIIα-expressing cells in the MPOA are responsible for the control of paternal behaviors. Moffitt and colleagues (2018) have identified at least 19 different neuronal phenotypes associated with Prlr expression in the MPOA. One potential candidate population of Prlr containing cells that may be important for paternal care are galanin (Gal) expressing cells, as Gal+ neurons have been shown to be critical for both maternal and paternal behaviors (Wu et al., 2014) and prolactin has been shown to excite Gal+ MPOA neurons (Stagkourakis et al., 2020). Although more work is needed to further characterize Prlr-expressing cells in the MPOA, we have identified that Prlr-expressing CaMKIIα forebrain cells are critical for the display of paternal behavior in fathers. While the MPOA population may be critical, it is unlikely that prolactin action on this circuit is limited to the MPOA, based on our c-fos data.

Since our data clearly showed that Prlr expression in the brain is critical for paternal behavior, our next question was at what time point(s) is prolactin exposure critical for the onset of paternal care? The mating-induced release of prolactin was previously thought to be involved in the refractory period between ejaculations in males (Brody and Krüger, 2006). However, a recent study has shown that this is not the case in mice (Valente et al., 2021), and therefore the function remains unclear. Since mating with ejaculation is the necessary stimulus for paternal care in mice, we hypothesized that mating-induced prolactin may serve as a signal to initiate the transition to paternal care. However, when prolactin was blocked with bromocriptine prior to mating, it did not have a significant effect on subsequent paternal behavior. In these studies, however, the control males appeared to perform less paternal behaviors relative to other control males (e.g. Fig 4). One factor that could account for this difference is that mated males in this study were not cohoused with pregnant females or pups in between mating and the pup retrieval test (as they were mated with ovariectomized females), whereas the other males tested (e.g. Fig 4) were co-housed with females and litters until day 3 postpartum (see Methods). Although paternal care in this mouse strain is not reliant on cohousing with a female, there can be potential additive effects of this, along with prior experience with the pups on paternal behavior (Brown, 1993). Notably, bromocriptine-treated animals did not show infanticidal responses, indicating that the mating-induced prolactin surge is also not required for the transition away from infanticide.

Aside from mating, basal circulating levels of prolactin are generally low and remain unchanged throughout the pup rearing period, which is maintained by TIDA neuron activity under normal physiological conditions (Stagkourakis et al., 2020). We also saw no acute increases in prolactin following pup interactions in control males. This pattern is in contrast to some other paternal rodent species such as Mongolian gerbils (Brown et al., 1995), Djungarian hamsters (Reburn and Wynne-Edwards, 1999), and California mice (Gubernick and Nelson, 1989) which show increased prolactin levels as fathers. Nonetheless, blocking pituitary prolactin secretion with bromocriptine resulted in similar pup retrieval deficits as observed in the Prlr CKC KO males, with prolactin administration following bromocriptine rescuing pup retrieval, suggesting the requirement for acute prolactin action during this behavior in fathers. In male mice, prolactin is the only identified lactogen to act through the Prlr, with growth hormone failing to bind to the Prlr in mice (Bartke and Kopchick, 2015) and placental lactogen only relevant in female mice during pregnancy (Cohick et al., 1996). It has also been confirmed that circulating prolactin can be transported past the blood brain barrier to access central Prlrs (Brown et al., 2016b; Barad et al., 2020), with bromocriptine treatment significantly eliminating pSTAT5 expression in the brain (Brown et al., 2010). Therefore, we can be confident that prolactin acting at the Prlr is mediating the observed effect on paternal behavior in these mice. It is important to note that this effect of prolactin promoting paternal behaviors is likely dependent on mating experience and/or other internal factors present in fathers, as injecting prolactin alone into virgin males did not cause them to show paternal behavior. Notably, blocking prolactin in sires did not return males to an infanticidal state; instead bromocriptine-treated males either ignored pups or displayed fewer paternal behaviors towards less pups.

In our studies, despite the clear deficits in pup retrieval behavior in both models of impaired prolactin action (Prlr CKC KO and bromocriptine-treated sires), a few manipulated males were also still able to exhibit paternal responses similar to control animals. A critical role for prolactin signaling on these circuits does not rule out regulatory roles for other hormones in supporting paternal behavior. Indeed, there is good evidence that hormones such as oxytocin, vasopressin, and estradiol also facilitate paternal responses (reviewed in Saltzman and Ziegler, 2014; Bales and Saltzman, 2016). Therefore, it is likely that other mechanisms, unaffected by our manipulations, were able to partially compensate for the loss of prolactin action in some of our animals. Despite these mitigating factors, however, the loss of prolactin action still had a profound effect on expression of paternal behavior, indicative of an important role for this specific hormonal cue in promoting pup retrieval, and likely other paternal behaviors, in mice.

Taken together, our data show that paternal behavior is dependent on basal levels of circulating prolactin acting at the Prlr on CaMKIIα-expressing forebrain neurons following mating experience. Our results also provide further support that the suppression of infanticide and the onset of paternal caring behaviors act via two different, parallel pathways and have separate neuroendocrine regulatory mechanisms (Kohl et al., 2017). To date, prolactin has been most well recognized for its role in lactation and the accompanying maternal care in female mammals. Contrary to its name, prolactin’s role in lactation may be a relatively recently-evolved adaptation within prolactin’s wider contribution to the regulation of mammalian parental care. These new data in male mice suggest that prolactin likely has a longstanding and conserved role in promoting the care of offspring, acting equivalently in both sexes. Given that humans are among the few mammalian species that exhibit paternal care (Dulac et al., 2014), it is likely that this mechanism will be important in men as well as women.

## METHODS AND MATERIALS

All procedures were approved by University of Otago Animal Ethics Committee.

## ANIMALS

Mice were housed in individually ventilated cages with shredded paper nesting material and kept in temperature-controlled rooms (22 ± 1°C) on 12:12 hour reverse light/dark cycles (lights on at 20:00 hrs) with *ad libitum* chow and water. All animals were tested during the dark cycle under sodium lighting (McLennan and Taylor-Jeffs, 2016). Mice were 8-12 weeks of age when used.

Adult C57BL/6J mice were sourced from the Taieri Resource Unit (University of Otago, Dunedin, New Zealand) from stock regularly refreshed from Jackson Laboratory (IMSR Cat# JAX:000664, RRID:IMSR_JAX:000664, The Jackson Laboratory, Bar Harbor, Maine, USA).

Generation of the Prlr-IRES-Cre knock-in mice has been previously described and characterized (Kokay et al., 2018; Aoki et al., 2019). Heterozygous Prlr-IRES-Cre mice were crossed with Ai9 Cre-dependent tdtomato reporter mice (B6.Cg-Gt(ROSA)26Sortm9(CAG-tdtomato)Hze/J; IMSR Cat# JAX:007909, RRID:IMSR_JAX:007909, The Jackson Laboratory, Bar Harbor, Maine, USA; Madisen et al., 2010), using methods previously described (Kokay et al., 2018) to generate Prlr-IRES- Cre/tdtomato reporter mice. As current immunohistochemistry methods lack the necessary sensitivity to reliably detect these low abundance Prlr in neurons, this reporter line is an invaluable tool for identifying Prlr-expressing cells in the brain.

Generation of Prlr^lox/lox^ mice has been previously described (Brown et al., 2016a). Animals with a prolactin receptor deletion in calcium/calmodulin-dependent protein kinase IIα (CaMKIIα)-expressing cells (Prlr^lox/lox^/CamK-Cre mice), GABAergic cells (Cre expression driven by the vesicular GABA transporter promoter; Prlr^lox/lox^/VGat-Cre mice), or glutamatergic neurons (Cre expression driven by the vesicular glutamate transporter 2 promoter; Prlr^lox/lox^/VGlut-Cre mice) were generated and genotyped as previously described (Vong et al., 2011; Brown et al., 2016a; Gustafson et al., 2020). CaMKIIα expression is predominantly restricted to the mouse forebrain, with no detection observed in the hindbrain (Solà et al., 1999) or outside the brain (Casanova et al., 2001), and is primarily expressed in excitatory cells, with the exception of the BST and cerebellar purkinje cells where CamKIIα and GABA are co-expressed (Benson et al., 1992; Liu and Murray, 2012). The pattern of recombination driven by this Cre-line has been extensively characterized elsewhere (Casanova et al., 2001; Brown et al., 2016a; Gustafson et al., 2020). Respective littermate cre-negative Prlr^lox/lox^ mice for each genotype were used as control animals. As previously described (Brown et al., 2016a), in the Prlr^lox/lox^ models above, the construct was designed such that the wild-type exon 5 and an inverted eGFP (Functional Enhanced Green Fluorescent Protein) reporter are flanked by lox66 and lox71 sites. As Cre-mediated inversion deletes the *Prlr* gene, eGFP is knocked into place so that eGFP can be used as marker for successful recombination, with control (cre-negative Prlr^lox/lox^) mice showing no eGFP expression (Brown et al., 2016a; Fig 3), indicative of intact *Prlr*.

## EXPERIMENTAL DESIGN

### Characterizing c-fos expression in Prlr-responsive neurons following pup interactions

Our first aim was to determine if Prlr-responsive neurons were activated in response to pup interactions in sires. To address this, we measured c-fos immunoreactivity (a marker of recent neural activation; protocol and image analysis described below) in Prlr-IRES-Cre-tdtomato reporter mouse sires that were randomly chosen to be either exposed to pups during a standard 30 min pup exposure assay (Fig 1A; described in detail below), or received no pup exposure (controls; Tachikawa et al., 2013; Tsuneoka et al., 2015). Males were left undisturbed in cages with pups for an additional 90 min following the test before brains were collected to enable maximal detection of pup-induced c-fos immunoreactivity (Tsuneoka et al., 2015). One pup-exposed male did not retrieve any pups and therefore was not included in the data analysis. C-fos, tdtomato (indicative of Prlr expression), and c-fos+tdtomato co-labeled immunoreactivity was quantified in the bed nucleus of the stria terminalis (BSTv), posteroventral division of the medial amygdaloid nucleus (MeApd), medial preoptic area (MPOA), and periventricular nucleus (PVN). These brain regions were chosen as they are known to be involved in paternal behavior (Tsuneoka et al., 2015; Bales and Saltzman, 2016) *and* express the Prlr (Kokay et al., 2018).

For comparison, we also exposed virgin Prlr-IRES-Cre/tdtomato reporter males to pups (Fig 2A; behavior test described below), to evaluate reproduction-driven changes in pup-induced neuronal activation. Following the pup test, males were left in the cage (without pups) for an additional 90 min following the pup exposure assay before brain collection. Finally, to confirm whether c-fos expression in Prlr-expressing neurons was unique to pup interactions, c-fos was assessed in a separate cohort of mice following mating. Once a male had ejaculated, the female was removed and brains were collected 90 min later. Additional control male brains were collected at equivalent times points to mated males.

### Effects of neuronal prolactin receptor knock-out on paternal behavior in sires

Following identification that populations of Prlr-expressing cells exhibited pup-induced c-fos immunoreactivity (Fig 1), we next wanted to test whether paternal behavior in mouse sires was dependent on neuronal expression of the prolactin receptor (Prlr). To do this, we generated 3 different conditional Prlr knockout lines (described above) in which Prlrs were genetically deleted from either glutamatergic (Prlr Vglut KO; Vong et al., 2011; Brown et al., 2017), GABAergic (Prlr Vgat KO; Vong et al., 2011; Brown et al., 2017), or from CaMKIIα expressing-forebrain neurons (Prlr CKC KO; Casanova et al., 2001; Brown et al., 2016a). Adult males from all 3 Prlr knock out lines were tested as sires for paternal behavior using the pup retrieval test (described below). The degree and distribution of Prlr deletion was assessed across the forebrain by eGFP immunoreactivity (described below), which is induced upon Cre-mediated recombination of the transgene (Brown et al., 2016a), with respective littermate Cre-negative Prlr^lox/lox^ (with intact *Prlr*) serving as control mice.

### Identifying critical periods of prolactin exposure for paternal behavior

Finally, we aimed to identify critical periods of prolactin exposure that may be important for both the transition and expression of paternal care. Using C57/B6J mice, we first characterized the circulating prolactin levels in males before, during, and after mating, and throughout the pup rearing period (procedures described below). Based on these findings, we first tested whether the mating-induced prolactin surge is required for the transition from the infanticidal (virgin) state to paternal (sire) state. C57BL/6J males were randomly assigned to be treated with either bromocriptine (a prolactin inhibitor, details below; n=9) or vehicle (n=9) 1.5 hours prior to mating, or bromocriptine 1.5 hours and ovine prolactin administered 45 mins prior to the mating test (n=7; prolactin injections mating behavior assay described below). Males had a blood sample taken prior to the bromocriptine/vehicle injection (baseline) and 30 min, 1 hr, 2 hr, 3 hr, and 4 hrs post-ejaculation to confirm treatment effects. Males remained singly housed after mating until they were tested 20-24 days later with foster pups (aged 3-5 days). Male lab mice do not differentiate between their own and foster pups and will readily retrieve either to the nest as sires (Alsina-Llanes and Olazábal, 2018). Males were blood sampled 1 hr prior to (pre-pup sample) and immediately following the pup exposure test (post-pup sample). For this pup retrieval test, males were presented with 2 foster pups. Although the pup retrieval tests generally uses 4 pups, we opted to use only 2 pups for these tests because we did not know if our manipulation would prevent the suppression of infanticidal behavior and we did not want to put additional pups at risk (Lonstein et al., 2002). Sufficient blood samples could not be collected at both time points for 1 control male and 3 bromocriptine treated male (Fig 6J). To confirm that bromocriptine had no effects on mating behavior, a separate group of C57BL/6J males were injected with bromocriptine (n=9) or vehicle (n=8) 1.5 hrs prior to the mating assay described above. Behavior was video recorded and scored as described below.

We next tested whether prolactin was required in sires to show paternal behaviors towards their pups. C57BL/6J males were mated and co-housed with the female until day 3 pp (see behavior assay). On the morning of testing (day 4 pp), males had a blood sample taken (pre-pup sample) and males were randomly assigned to be treated with either bromocriptine (n=8) or vehicle (n=6) 1.5 hrs prior to the pup test, or bromocriptine 1.5 hours and ovine prolactin 45 mins prior to the pup test (n=6). Four pups (age 4 days) from the male’s original home cage were used as pup stimuli for the pup exposure assay. A second blood sample was taken immediately following the 30 min pup exposure test (post-pup sample). A sufficient amount of blood could not be collected from 1 control male and 1 bromocriptine-treated male, hence only 5/6 control males and 7/8 bromocriptine males had blood samples measured (Fig 4Q).

Finally, to test whether prolactin can induce paternal care in virgins, C57BL/6J males were treated with prolactin or vehicle before the pup exposure assay. Males were randomly assigned to be injected either 45 minutes prior to the pup test (Fig S4A-C: acute effects of prolactin; n=7 per group) or 3 weeks prior to the pup test (FigS4D-F: delayed effects of prolactin, n=10 per group). To verify that prolactin is not required for infanticidal behavior, virgin C57BL/6J males were either injected with bromocriptine (as described above, n=9) or vehicle (n=9) for 1.5 hrs before the virgin pup test commenced, as described below.

## BEHAVIOR ASSAYS AND QUANTIFICATION

### Sire pup exposure assay

The pup exposure assay used in sires (Figs 1, 4, and 6) was based on the pup assay used in Tsuneoka et al., 2015. Briefly, virgin males were paired with a novel wild-type female and co-housed together until day 3 post-partum (pp; the first day pups were observed was counted as day 1 pp). On day 3 pp, males were removed and singly housed in a new cage for 24 hrs. Males were separated from their litters in order to eliminate any potential c-fos expression from previous interactions with pups in the home cage (for animals used in Fig 1) and to elicit a maximal paternal response upon reunion with pups (Tsuneoka et al., 2015). On the following day (day 4 pp), males were brought into a quiet testing room and allowed to acclimate for 15 min. Cage lids were replaced with clear plexiglass and a video camera was placed directly above the cage to record behaviors. Following the acclimation period, four pups (age 4 days) from the male’s original home cage were placed in his new cage, opposite his nest. Paternal behaviors were recorded for 30 min. Control males (in Fig 1) underwent all procedures as pup-exposed males except that only a hand was briefly placed in the cage, but no pups were added. All pup retrieval testing was conducted between 09:00 hrs and 12:00 hrs.

For all pup exposure tests, the behaviors measured included sniffing pups (nose in contact with part of a pup’s body), retrieving pups (picking up pups with their mouth and carrying into the nest), nesting alone (in nest with no pups), or huddling (hovering over at least one pup in the nest). Behaviors were scored from videos using the program BORIS (Friard and Gamba, 2016) using a scan-sampling method every 15 seconds (Lonstein and Fleming, 2001; Tsuneoka et al., 2015). This method yields similar proportions of time spent engaged in each behavior as when full durations of each behavior are recorded (unpublished data). The number of instances for each behavior was divided by the total number of observations per video (e.g., 120 observations for a 30 min video) to give the proportion of time an animal engaged in each behavior. The researcher scoring videos were blind to conditions.

### Virgin pup exposure assay

For pup tests using virgin males (Fig 2, S3), males were separated into individual cages 24 hrs before testing. On the day of testing, two foster pups (aged 3-6 days old) were placed in the cage, opposite of the nest, and behavior was observed for 10 min. All virgin pup tests were observed live and if males attacked a pup, the pups were immediately removed, euthanized, and testing ceased. If males ignored pups, then pups were removed after the 10 min test and returned to their home cage. The latency to attack or retrieve pups was recorded, as well as behavioral outcomes at the end of the test (attack, ignore, or paternal). Virgin control males (in Fig 2) underwent the all similar procedures, except only a hand was briefly placed in the cage, but no pups were given. All virgin behavioral testing took place between 09:00 and 12:00 hrs.

### Mating behavior assay

Males were singly housed and tested in their home cage. For mating tests, one sexually receptive female was placed in the male’s home cage and mating behavior was both observed live and video recorded. Once a male had ejaculated, testing ceased, and the female was returned to her original home cage. Female stimuli used for all mating tests were reproductively-experienced, ovariectomized wildtype animals which were brought into receptivity by using a standard protocol of injecting estradiol (0.01mg injection/sc, dissolved in sesame oil, vol=0.1ml, Sigma 815) 48 hrs prior and progesterone (0.05mg injection/sc, dissolved in sesame oil, vol=0.1ml, Sigma P0130) 4 hrs before testing (Liu et al., 2020). All females were injected at 09:00 hrs and mating behavior tests took place between 13:00-17:00 hrs.

Mating behaviors were scored from videos using the program BORIS (Friard and Gamba, 2016), including latency to first mount and to ejaculate, the number of mounts and intromissions, and duration of each mounting bout. The researcher scoring videos was blind to condition.

## BLOOD SAMPLING AND PROLACTIN ASSAY

### Blood sampling procedure

Whole blood samples were collected from the tail vein of mice following methods described in Steyn et al., 2011; Guillou et al., 2015. Mice were handled daily in a gentle restraint device (cardboard tube) for 3 weeks prior to blood collection to habituate mice to the procedure and to reduce stress at the time of blood collection. At the beginning of the sampling period, the tail tip (<1mm from the end of the tail) was cut with a sharp scalpel blade, and then the tail was gently squeezed to encourage a drop of blood to form at the site of the cut. Twelve μL of whole blood were collected with a pipette at each sampling point and immediately diluted 1:10 in 0.01M phosphate buffered saline containing 0.05% Tween and 0.2% bovine serum (PBST-BSA) and snap-frozen on dry ice. Samples were stored at - 80°C until analysis.

### Blood sampling before and after mating

Data presented in Fig 6A and C are from the same cohort of C57/B6J mice (n=12). These males were housed in a normal 12:12 light cycle room (lights on at 08:00 hrs) and were blood sampled as virgins at 09:00 hrs (light phase) and 21:00 hrs (dark phase) on the same day. Males were then paired with a wildtype female. Mating and pregnancies were monitored by daily morning checks for vaginal plugs and female weight gain. Each male was blood sampled between 09:00-10:00 hrs on days 12, 20, 35, 50, and 60 after mating (day of plug was determined as day 1). Males remained with females and pups during the duration of blood sampling period. The number of missing blood samples (due to inability to collect sufficient quantities of blood) were as follows: day 12 (n=3), day 20 (n=1), day 50 (n=1), and day 60 (n=1).

### Blood sampling during mating

For blood data collected during mating (Fig 6B and S2A), all animals were initially group-housed and habituated to handling in a restraint device (paper towel roll) daily for 3 weeks prior to blood sampling. Males were separated into individual cages two days before testing. Approximately 1.5 hrs before the test started, a baseline blood sample was taken, followed by an injection of either bromocriptine (n=9) or vehicle (n=7) (injections described below; bromocriptine and vehicle data shown in Fig S2A; vehicle data only shown in Fig 4B). A receptive female was added to the cage and mating behavior was observed. Once a male had ejaculated, a blood sample was collected immediately and the female was returned to her home cage. Further blood samples were collected 30 min, 1 hr, 2 hr, 3 hr, and 4 hrs post-ejaculation.

### Prolactin ELISA

Blood prolactin concentrations were measured with an enzyme-linked immunosorbent assay (ELISA), which has been described elsewhere (Kirk et al., 2017). Values that were not detectable by the ELISA were assigned a value of 0.1 ng/mL (the limit of detection; range 0.1-20 ng/mL). Blood samples in Fig 6A and 6C were run on 1 plate and blood samples in Fig 6B and S2A were run across 4 plates. The Inter-plate CV was 2.6% and the intra-plate CVs were 0.23 - 2.22%.

## DRUG TREATMENTS

Prolactin secretion was blocked by bromocriptine (200μg injections/sc, vol= 0.3ml, Sigma B2134; vehicle = 10% ethanol dissolved in sterile saline). Bromocriptine is a D2 agonist which prevents prolactin secretion from the pituitary within one hour (Brown et al., 2010; Valente et al., 2021). Circulating prolactin can be transported past the blood brain barrier to access central Prlrs (Brown et al., 2016b), which is prevented by bromocriptine treatment (Brown et al., 2010). Ovine prolactin was used (Fig 5, 6, S3) at 5 mg/kg injection/ip, dissolved with 10% ethanol in saline, Sigma L6520 (Brown et al., 2010).

## BRAIN COLLECTION, IMMUNOHISTOCHEMISTRY, AND IMAGE ANALYSIS

### Brain collection

To collect brains for immunohistochemistry (IHC), mice were deeply anaesthetized with sodium pentobarbital (100 mg/ kg^-1^; ip injection) before transcardial perfusion with 4% paraformaldehyde in 0.1 mol L^-1^ phosphate buffer (pH 7.4). Brains were post-fixed in the same fixative overnight before being cryoprotected in 30% sucrose solution for 2 days and stored at -80°C. Brains were cut into 3 series of 30 μm-thick sections on a freezing microtome and kept in cryoprotectant at -20° C until processing for IHC.

### C-fos immunofluorescence

The protocol used was adapted from Brown et al., 2019. Briefly, sections were incubated in rabbit anti-cfos primary antibody (rabbit polyclonal Anti-c-fos, 1:5000, Abcam Cat# ab190289, RRID:AB_2737414, Abcam, Melbourne, Australia) for 48 hours at 4°C, followed by a 60 min incubation in biotinylated goat anti-rabbit IgG (1:200; Vector Laboratories Cat# BA-1000, RRID:AB_2313606, Vector Laboratories, Peterborough, United Kingdom). Sections were then incubated in Vector Elite avidinbiotin-horseradish peroxidase complex (1:100) for 45 min, before a 20 min incubation in Biotin-XX Tyramide (0.3%; Invitrogen, Carlsbad, CA). Finally, sections were incubated in a Streptavidin 647 IgG (1:400; AlexaFluor; Invitrogen) for 2 hrs at 37°C. Images were captured using a Nikon A1 inverted confocal using a 20X objective. Z-stacks were collected with images taken 1.4 μm apart. For all IHC runs (including below), control sections that had the primary antibody omitted were included in each IHC batch. No specific staining was observed in any of these sections.

### C-fos image analysis

The number of tdtomato labeled cells, c-fos labeled cells, and double-labeled cells (with both tdtomato and c-fos) were manually counted in each region of interest using NIS-Elements AR Analysis software, Nikon (NIS-Elements, RRID:SCR_014329) by a researcher blind to conditions. Cell count numbers were divided by the area analyzed (mm^2^) to calculate the density of cells per region for each animal. Area outlines were drawn based on the stereotaxic mouse brain atlas (Franklin and Paxinos, 2013). One hemisphere per section for each brain region was counted for each animal. Images of compressed z-stacks were pseudo-colored magenta and cyan in order to be color-blind friendly and prepared in FIJI distribution of image J (National Institutes of Health, Bethesa, MD; RRID:SCR_002285).

### Chromogen eGFP IHC

To quantify eGFP immunoreactivity across our 3 Prlr^lox/lox^ models (Fig 3), separate groups of male Prlr^lox/lox^/CamK-Cre+, Prlr^lox/lox^/VGlut-Cre+, Prlr^lox/lox^/VGat-Cre+ and respective littermate Prlr^lox/lox^Cre-controls (n=6 per group) mice were perfused and brains were collected (as described above) at 8-12 weeks of age. The IHC protocol was performed as described in Brown et al., 2016a and Kokay et al., 2018 using rabbit polyclonal anti-GFP antibody (1:30,000, Molecular Probes Cat# A-6455, RRID:AB_221570, Invitrogen-ThermoScientific, United States) and biotinylated goat anti-rabbit IgG secondary antibody at 1:200 (Vector Laboratories Cat# BA-1000, RRID:AB_2313606, Vector Laboratories, Peterborough, United Kingdom).

### Chromogen pSTAT5 IHC

To measure pSTAT5 activity in Prlr^lox/lox^/CamK-Cre mice (Fig 5), a separate cohort of Prlr^lox/lox^/CamK-Cre+ and Prlr^lox/lox^Cre-control male (n = 6 per group) were injected with ovine prolactin (as described above) 45 min before perfusion, as in Brown et al., 2010. Immunhistochemcial labelling for pSTAT5 was undertaken as previously described (Brown et al., 2010) using Phospho-Stat5 (Tyr694) primary antibody (pSTAT5 Tyr 694, 1:1,000; Cell Signaling Technology Cat# 9351, RRID:AB_2315225) and biotinylated goat anti-rabbit IgG secondary antibody at 1:200 (Vector Laboratories Cat# BA-1000, RRID:AB_2313606, Vector Laboratories, Peterborough, United Kingdom).

### Chromogen image analysis

Images were taken on an Olympus AX70 brightfield microscope using 4X and 10X objectives. One hemisphere per section for each brain area was counted for each animal. Brain regions of interest were drawn based on the stereotaxic mouse brain atlas (Franklin and Paxinos, 2013). For all cell counts, the optical densities in each section were thresholded and automatically quantified in Image J software (National Institutes of Health, Bethesa, MD; RRID:SCR_003070) and expressed as cell density (number of counted cells per mm^2^ of the area measured). Image analysis was performed by a researcher blind to conditions.

## STATISTICAL ANALYSIS

No statistical methods were used to predetermine sample sizes but our sample sizes are similar to those reported in previous publications (e.g.,Wu et al., 2014; Brown et al., 2017). Data was analysed using PRISM 8 and 9 (GraphPad Prism, RRID:SCR_002798, San Diego, CA USA). In all cases, data was checked for normality, all tests were two-tailed, and significance was accepted if p-values were less than 0.05. All statistical analyses, including group means/medians, standard error of the mean (SEM), 95% confidence intervals (CI), and R squared (eta squared) values for effect size estimates are provided in Tables 1-6 and S1-S3, with the corresponding figure panels listed.

Analyses between 2 groups (control and treated/knock-out groups) were conducted with t-tests if data were normally distributed or Mann-Whitney U tests if data were not normally distributed (Figs 1-6; S1-S3; see Tables 1-6; S1-S3).

Group differences in eGFP-labelled cell densities between the 3 Prlr knockout models (Prlr CKC, Vglut, and Vgat KO males; Fig 3) were analyzed using a one-way ANOVA. As Cre-negative Prlr^lox/lox^ mice (with intact *Prlr*) show no eGFP immunoreactivity in the brain (Fig 3A-F), eGFP was only quantified and compared in Cre+ males. Pairwise comparisons between models were conducted using Tukey’s multiple comparisons test. Differences in eGFP-labelled cell densities between the 3 knockout models were analyzed for each brain region separately (Table 3).

Group differences in the latency to retrieve pups (Fig 4D,J,P; Fig 6D,K; Fig S3C,F), latency to attack pups (Fig S3B,E,H), latency to first mount a female (Fig S2B), and latency to first ejaculate (Fig S2C) were analyzed using survival analysis and curve comparison with the Mantel-Cox Log-rank test as in Swart et al., 2021. Hazard ratios (Mantel-Haenszel tests) and 95% confidence intervals are reported as an estimate of the effect size for the comparison of latency data.

Differences in blood prolactin concentrations between the light and dark phase in virgin males (Fig 6A) and pre- and post-pup test (Fig 6J,Q) were analyzed using a paired t-test. Blood prolactin concentrations across the time post-mating (Fig 6B) were analyzed using a repeated measures ANOVA. Post-hoc comparisons between groups (time of sample post-mating) were compared to the baseline sample group (pre-mating) using Dunnett’s test to correct for multiple comparisons. Blood prolactin concentrations across the days post-mating (Fig 6C) were analyzed using a repeated measures ANOVA. Post-hoc comparisons between groups (day of sample post-mating) were analyzed using Tukey’s test to correct for multiple comparisons. Differences in blood prolactin concentrations between bromocriptine and vehicle control groups following mating (Fig S2A) were analyzed using a 2-way repeated measures ANOVA, with post-hoc comparisons made using Bonferroni’s test to correct for multiple comparisons. Differences in paternal behavior between control, bromocriptine, and bromocriptine+prl groups (Fig 6 E-I, L-P) were analyzed using a One-way ANOVA if data were normally distributed or a Kruskal-Wallis test if data were not normally distributed (see Table 6). Post-hoc comparisons between the treated groups were compared to the vehicle control group using Dunnett’s multiple comparisons test (ANOVA tests) or Dunn’s multiple comparisons test (Kruskal-Wallis tests).

The proportion of virgin males showing attacking, ignore, or paternal responses was compared between treated males (acute prolactin, delayed prolactin, or bromocriptine) and their respective vehicle-treated controls using two proportion z-tests (Fig S3A,D,G).

## ACKNOWLEDGEMENTS

This work was support by the Health Research Council of New Zealand, the Marsden Fund - Royal Society of New Zealand, and a Project Support Grant from the British Society of Neuroendocrinology. We are grateful for the work of Pene Knowles who genotyped the animals and Genieve Yeo for assistance with behavior video coding.

## AUTHOR CONTRIBUTIONS

KOS: Conceptualization, Methodology, Validation, Formal analysis, Investigation, Writing – Original Draft, Visualization, Funding acquisition. RSEB: Conceptualization, Methodology, Validation, Writing - Review & Editing, Supervision. DRG: Conceptualization, Methodology, Resources, Writing - Review & Editing, Supervision, Funding acquisition.

## SUPPLEMENTRY MATERIALS

**Figure S1.**
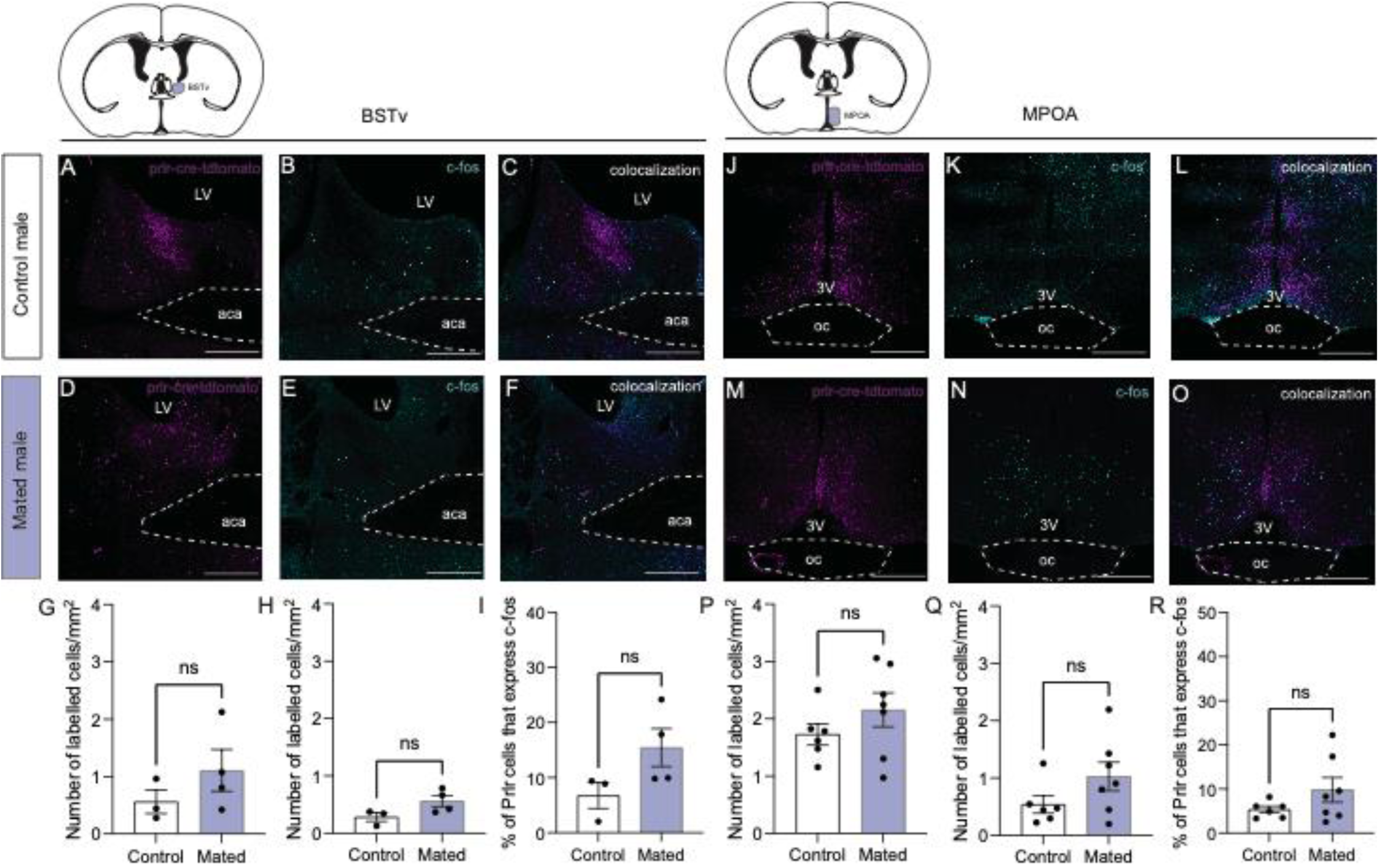
Prolactin-receptor containing neurons are not activated during mating in the BSTv or MPOA. C-fos was assessed in the bed nucleus of the stria terminalis (BSTv) and medial preoptic area (MPOA) of Prlr-IRES-Cre-tdtomato reporter males following their first mating experience compared to control males who were not exposed to a female. (A-O) Representative images from control males (top row) and mated males (bottom row) of tdtomato-labelled cells (magenta, indicative of a Prlr-expressing cell), c-fos (cyan), and the number of colocalized (tdtomato + c-fos, white). Note that there were no differences in the density of tdtomato-balled cells (G, P), c-fos-labeled cells (H, Q), or the percentage of Prlr cells expressing c-fos in the BSTv or MPOA (I,R) between controls (white bars) and mated males (purple bars). Bar graphs show individual data points (black circles) and mean ± SEM; ns = non-significant (p>0.05). LV= lateral ventricle, 3V = third ventricle, aca = anterior commissure, oc = optic chiasm, opt = optic tract. Scale bars = 50 μm.

**Figure S2.**
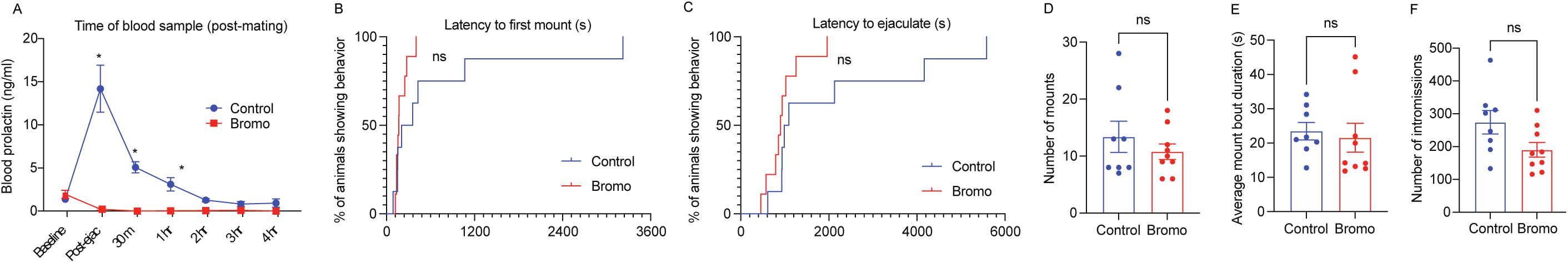
Suppressing prolactin at mating does not affect mating behavior. Mating behavior was assessed in C57/BL6J male mice that were treated with bromocriptine (n=9, red) or vehicle (n=7, blue) 1.5 hours prior to mating. (A) Vehicle-control males showed the expected transient rise in prolactin following ejaculation, whereas bromocriptine suppressed prolactin for up to 4 hrs post-mating. Single asterisks indicate a significant difference between control and bromocriptine-treated groups (p<0.05). Bromocriptine did not affect the latency to first mount (B), latency to ejaculation (C), the number of mounts (D), mount duration (E), or number of intromission (F) compared to control males.

**Figure S3.**
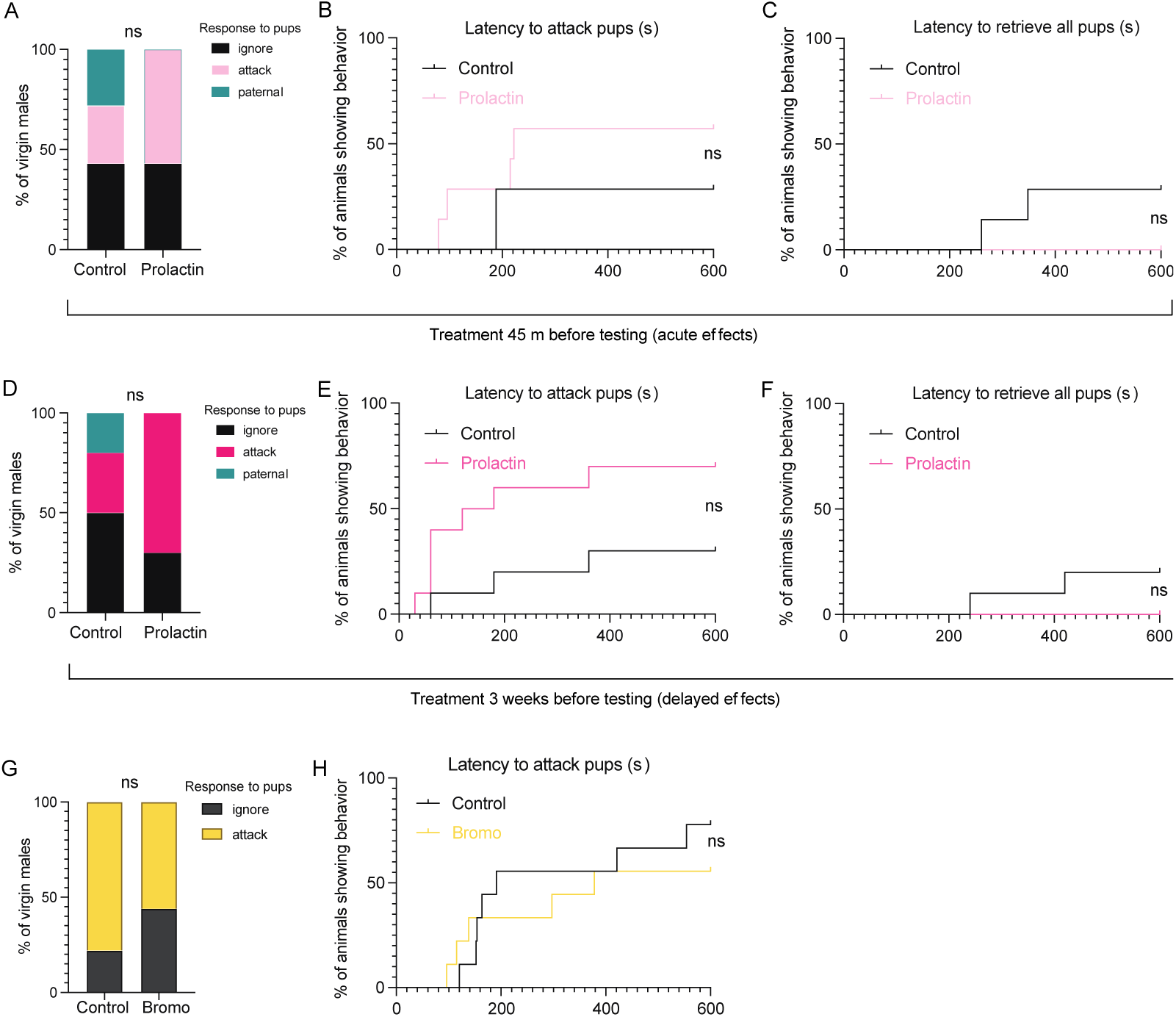
Prolactin exposure does not affect attacking or paternal behavior in virgin male mice. Neither acute prolactin treatment (45 min prior to the pup retrieval test) or delayed prolactin exposure (administered 3 weeks prior to the pup test) in virgin C57/B6J mice affected the proportion of males which showed attacking, ignoring, or paternal responses towards pups (A,D), the latency to attack (B,E), or latency to retrieve pups (C,F) compared to vehicle-control males. Bromocriptine treatment (1.5 hours prior to the pup retrieval test) also had no effect on the proportion of males which showed attacking or ignoring responses towards pups (G), or the latency to attack pups (H). Note that none of the virgin males showed paternal responses in this experiment. (s)=seconds; ns = non-significant (p>0.05).

**Table S1.** Statistical analysis for Figure S1

| Corresponding Figure | Corresponding Panel | Analysis |  |  |  |  |  |  |  |  |  |  |
| --- | --- | --- | --- | --- | --- | --- | --- | --- | --- | --- | --- | --- |
| Figure S1 |  |  |  |  |  |  |  |  |  |  |  |  |
|  |  | Test | Brain Region | Parameter | t | df | p-value | Mean - Control group | Mean - Mated group | SEM | 95% confidence interval | R squared (eta squared) |
|  | G | t-test | BSTv | tdTomato (Number of labelled cells/mm2) | 1.18 | 5.00 | 0.293 | 0.56 | 1.11 | 0.46 | -0.649 to 1.740 | 0.22 |
|  | H | t-test | BSTv | C-fos (Number of labelled cells/mm2) | 2.08 | 5.00 | 0.092 | 0.28 | 0.56 | 0.13 | -0.065 to 0.623 | 0.46 |
|  | I | t-test | BSTv | % of Prlr cells that express C-fos | 1.91 | 5.00 | 0.115 | 6.76 | 15.45 | 4.55 | -3.020 to 20.390 | 0.42 |
|  | P | t-test | MPOA | tdTomato (Number of labelled cells/mm2) | 1.18 | 11.00 | 0.264 | 1.73 | 2.16 | 0.36 | -0.372 to 1.226 | 0.11 |
|  | Q | t-test | MPOA | C-fos (Number of labelled cells/mm2) | 1.60 | 11.00 | 0.138 | 0.54 | 1.03 | 0.31 | -0.185 to 1.162 | 0.19 |
|  | R | t-test | MPOA | % of Prlr cells that express C-fos | 1.43 | 11.00 | 0.180 | 5.33 | 9.79 | 3.11 | -2.391 to 11.310 | 0.16 |

**Table S2.**
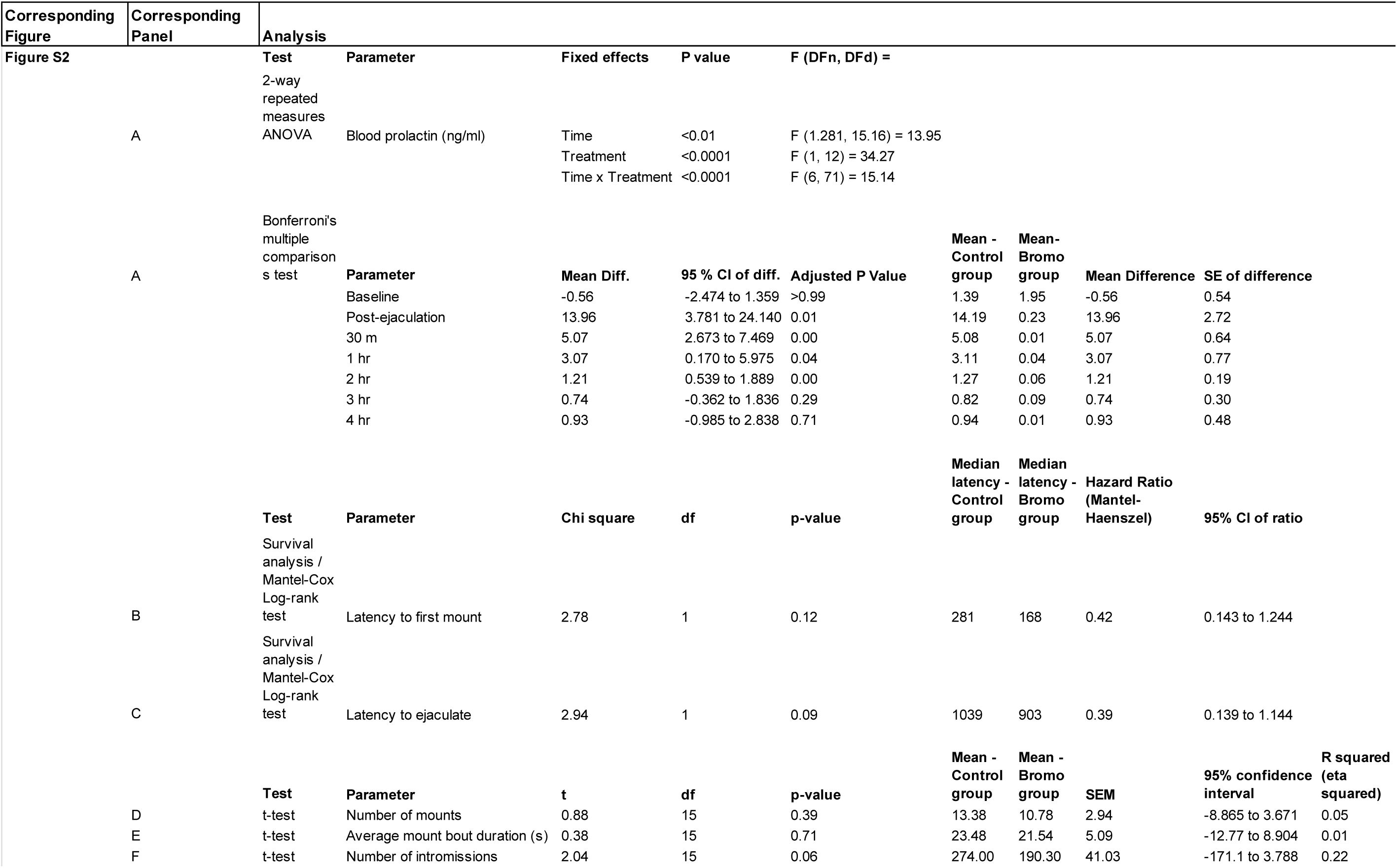
Statistical analysis for Figure S2

**Table S3.**
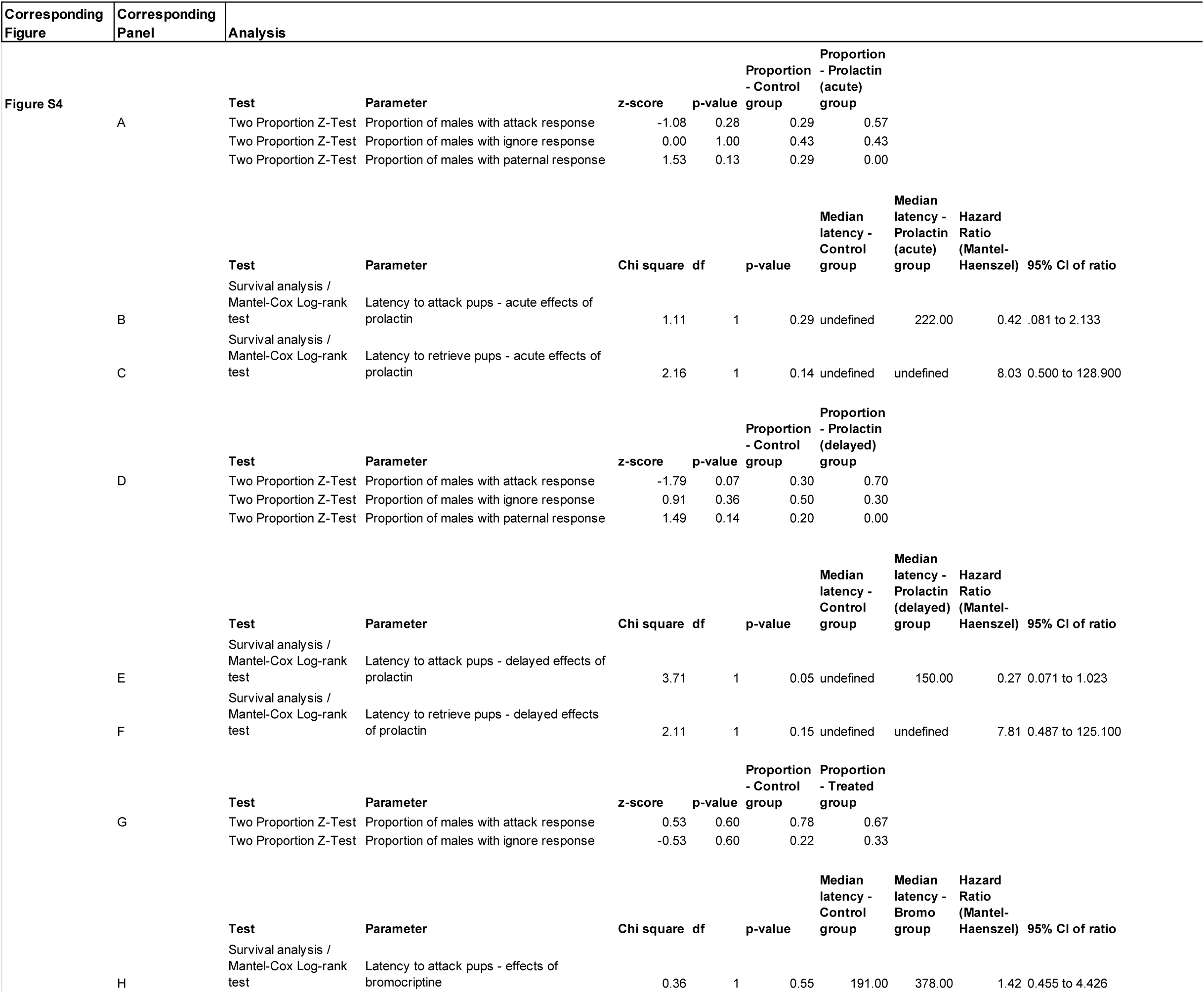
Statistical analysis for Figure S3

**Movie S1. Prolactin receptor knockout in CaMKIIα-expressing neurons disrupts up retrieval behavior (separate file).** Video shows a representative example of a control sire (cre-negative Prlr^lox/lox^) and a CaMKIIα-neuron-specific prolactin receptor knock-out (Prlr CKC KO) sire behavior during the pup retrieval test. Each video shows the first two minutes of the test and is sped up to 4X speed. Note that the control male quickly retrieves pups to the nest. In contrast, while the Prlr CKC KO male investigates/sniffs the pups, he does not retrieve the pups to the nest.

